# miR-467 regulates inflammation and blood insulin and glucose

**DOI:** 10.1101/666545

**Authors:** Jasmine Gajeton, Irene Krukovets, Revanth Yendamuri, Dmitriy Verbovetskiy, Amit Vasanji, Lidiya Sul, Olga Stenina-Adognravi

## Abstract

Obesity is associated with inflammation and insulin resistance (IR), but the regulation of insulin sensitivity (IS) and connections between IS and inflammation remain unclear. We investigated the role of miR-467a-5p, a miRNA induced by hyperglycemia, in regulating inflammation and blood glucose handling.

We previously demonstrated that miR-467a-5p is induced by hyperglycemia and inhibits the production of thrombospondin-1 (TSP-1), a protein implicated in regulating inflammation. To investigate the role of miR-467 in blood glucose handling and tissue inflammation, WT C57/BL6 mice were fed chow or Western diet from 5 to 32 weeks of age and injected weekly with miR-467a-5p antagonist. Inhibiting miR-467a-5p resulted in 47% increase in macrophage infiltration and increased *Il6* levels in adipose tissue, higher plasma insulin levels (98 vs 63 ng/mL), and 17% decrease in glucose clearance without increase in weight or HDL/LDL. The antagonist effect was lost in mice on Western diet. Mice lacking TSP-1 lost some but not all of the miR-467 effects, suggesting *Thbs1^−/−^* (and other unknown transcripts) are targeted by miR-467 to regulate inflammation.

miR-467a-5p provides a physiological feedback when blood glucose is elevated to avoid inflammation and increased blood glucose and insulin levels, which may prevent IR.

## 1. INTRODUCTION

Genome-wide analyses have uncovered important roles of microRNAs in the pathogenesis of diabetes mellitus ^1^, including evidence to suggest a tight regulation between microRNAs, glucose metabolism, and inflammation ^2–4^. The expression of microRNAs can be further altered by a variety of stressors, e.g., changes in blood glucose or pro-inflammatory cytokines ^5^; regulation of miRNAs adds another layer of complexity in regulating targets. Diet-induced obesity and increased blood glucose levels correlate with chronic inflammation and development of IR ^6–14^. However, the sequence and causality of pathological changes leading to IR, including naturally occurring feedback mechanisms preventing the transition to IR in response to elevated blood glucose, are poorly understood.

Macrophage infiltration in adipose tissue is thought to be a main contributor in promoting chronic inflammation and development of IR. Islet inflammation promotes impaired β-cell function and subsequent failure, which occurs before the onset of type 2 diabetes (T2D) ^15–17^. Thrombospondin-1 is an extracellular matrix protein involved in regulation of tissue remodeling and inflammation. Studies in *Thbs1*^−/−^ mice suggest that a lack of TSP-1 may alleviate macrophage accumulation and the pro-inflammatory phenotype observed in insulin resistant metabolic organs, thus protecting the animals from diet-induced inflammation and IR ^18–20^.

We recently reported that miR-467a-5p is rapidly upregulated by high glucose *in vitro* and *in vivo* and regulates angiogenesis by targeting *Thbs1* mRNA ^21–24^. Others report this miRNA prevents vascular inflammation by targeting Lipoprotein Lipase in macrophages ^21,22,25–28^. Yet, the physiological function of miR-467a-5p and the physiological significance of its rapid upregulation by hyperglycemia remained unknown. In this work, the effects of a miR-467 antagonist on blood glucose and insulin levels and inflammation in adipose tissue and pancreas were examined in wild type (WT) and *Thbs1^−/−^* mice to understand the role of miR-467a-5p and its target, TSP-1, in regulating inflammation in tissues and in blood glucose handling.

## 2. MATERIALS AND METHODS

Detailed description of methods is provided in the Online Supplement.

### 2.1 Experimental animals

Animal procedures were approved by IACUC. Male WT C57BL6 (n=10/group) or *Thbs1^−/−^* (n=7/group) mice were fed a chow or Western diet (TD.88137, 40-45% kcal from fat, 34% sucrose by weight, Envigo) starting at 4 weeks of age and injected weekly with a miR-467a-5p antagonist (2.5 mg/kg body weight) (or a control oligonucleotide with no predicted targets in mouse or human genomes ^22,29^), intraperitoneally, starting at 5 weeks of age until the end of the experiment.

### 2.2 miR-467a-5p mimic and the miR-467a-5p antagonist

The miR-467a-5p mimic and the control oligonucleotide were purchased from Dharmacon. The custom LNA-modified miR-467a-5p antagonist and a control oligonucleotide were from Qiagen.

### 2.3 Glucose and insulin tolerance tests (GTT and ITT)

Glucose (2 g/kg body weight) or insulin (50 µg/kg) (Sigma) were injected intraperitoneally. Blood glucose levels were measured 0 – 180 min after injections using an AlphaTRAK glucometer.

### 2.4 Induction of diabetes in mice

Male mice were injected intraperitoneally with streptozotocin (STZ, 50 mg/kg, Sigma) for 5 consecutive days. Mice with blood glucose >250 mg/dL were selected for experiments.

### 2.5 Blood cell counts, HDL/LDL cholesterol, and cytokines in blood

Blood was collected by cardiac puncture and circulating blood cell counts were analyzed using an ADVIA 120 Hematology System (Siemens). Plasma insulin was measured using Insulin Mouse ELISA kit (Thermo).

A custom U-plex Assay Platform (MSD) was used to assess plasma levels of CCL2 (MCP-1), IL-10, CXCL1, and VEGF-A.

HDL and LDL cholesterol were measured using the HDL and LDL/VLDL quantification kit (BioVision) at end of the experiment.

### 2.6 Immunohistochemical staining

Visceral (omental) adipose tissue and pancreas were fixed in 4% formaldehyde (Electron Microscopy Sciences) for 24 hours and stained using VECTASTAIN ABC-HRP Kit (Vector Labs) with corresponding primary antibodies. Slides were scanned using Leica SCN400 or Aperio AT2 at 20X magnification. Quantification of positive staining was performed using Photoshop CS2 (Adobe) or Image Pro Plus (7.0).

### 2.7 Cell culture

RAW264.7, THP-1, βTC6 and 3T3-L1 cells were purchased from ATCC and cultured according to ATCC directions.

### 2.8 Isolation of bone marrow-derived macrophages (BMDM)

was performed as described in ^30^.

### 2.9 Glucose stimulation of RAW264.7, differentiated THP-1, βTC6, and BMDM

Cells were stimulated with 30 mM D-glucose High Glucose, “HG” (Sigma) for 6 hours (RAW 264.7 and BMDM), 3 hours (3T3-L1) or 30 minutes (βTC6).

### 2.10 Transfection of cultured cells

Transfections were aided with Oligofectamine (Invitrogen).

### 2.11 Oil Red O Staining

Differentiated 3T3-L1 cells were stained in the Oil Red O solution for 10’ at RT.

### 2.12 RNA Extraction and RT-qPCR

RNA was isolated using Trizol reagent (Thermo).

RNA was polyadenylated using NCode miRNA First-Strand cDNA Synthesis kit (Invitrogen) or miRNA 1st strand cDNA synthesis kit (Agilent). Real-time qPCR amplification was performed using SYBR GreenER™ qPCR SuperMix Universal (Thermo) or miRNA QPCR Master Mix (Agilent).

To measure expression of inflammatory markers, Real-time qPCR was performed using TaqMan primers for *Tnf*, *Il6*, *Ccl2*, *Il1b*, *Il10*, *Ccl4, Cd68, Slc2a1, Slc2a2, Slc2a4, G6pc, Fbp1* (Thermo) and TaqMan Fast Advanced Master Mix (Thermo) as described ^31^.

β-actin primers (CAT GTA CGT TGC TAT CCA GGC, IDT) were used for normalization by the the 2^−ΔΔCt^ method. All samples were assayed in triplicates using a fluorescence-based, real-time detection method (BioRad MyIQ RT-PCR, Thermo).

### 2.13 Statistical analysis

Data are expressed as the mean value ± S.E.M. Statistical analysis was performed using GraphPad Prism 5 Software. Student’s t-test and ANOVA were used to determine the significance of parametric data, and Mann-Whitney test was used for nonparametric data.

## 3. RESULTS

### 3.1 Injections of miR-467a-5p antagonist increase macrophage accumulation in adipose tissue and pancreas

Inflammation and macrophage infiltration in tissues are associated with, and often precede, IR ^32–37^. Macrophage accumulation in adipose tissue and pancreas was assessed by immunohistochemistry using an anti-MOMA-2 or anti-CD68 antibody (Figures 1A – 1D; Figures S1A – S1D show H&E and images of immunohistochemical staining).

**Figure 1.**
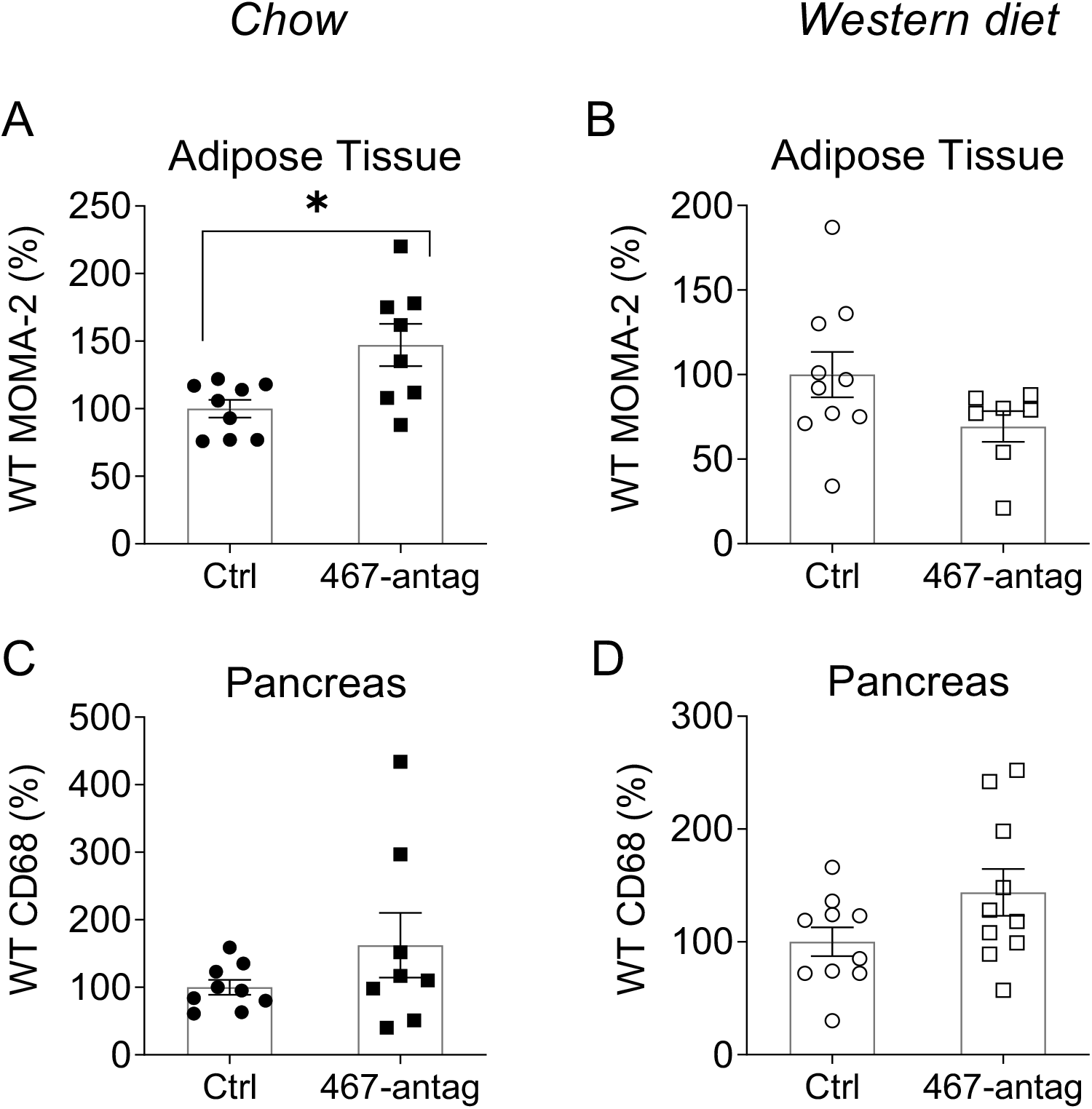
miR-467 antagonist increases macrophage accumulation in adipose tissue and pancreas from WT chow-fed mice. Macrophage accumulation in adipose tissue from WT mice on chow diet (A) and Western diet (B) was determined by anti-MOMA-2 staining. Positive staining was normalized to mean adipocyte area for adipose tissue since adipocyte sizes were changed between groups. Macrophage accumulation in pancreas from WT mice on chow diet (C) and Western diet (D) was determined by anti-CD68 staining. Data are relative to ctrl oligo. n=10 mice/group. \**P*<.05

Male WT mice on chow diet were injected weekly with a miR-467a-5p antagonist (2.5 mg/kg) for 32 weeks, starting at 5 weeks of age. Injections of the miR-467a-5p antagonist increased macrophage accumulation in AT of chow-fed mice by 47% (Figure 1A). Baseline AT macrophage infiltration was increased in mice on Western diet (65.4%, Figure S2A) without further increase in response to antagonist (Figure 1B).

In pancreas, macrophage infiltration tended to increase in antagonist-injected mice on either diet (62.4% increase, chow-fed and 43.9% increase, Western diet (Figures 1C, 1D) and was significantly increased by the Western diet (73.3%, Figure S2B).

Changes in tissue macrophage infiltration in response to the antagonist were not explained by the number of monocytes in blood (Figure S3): blood monocyte numbers were not increased by the antagonist. The Western diet increased numbers of circulating monocytes and white blood cells (Figures S3A – C).

### 3.2 miR-467a-5p antagonist has differential effects on the expression of inflammatory markers in adipose tissue

Adipose tissue expression of *Il6, Tnf*, *Ccl2*, *Ccl4*, or *Il1b* was assessed by RT-qPCR in chow or Western diet-fed WT mice (Figures 2A, 2B, respectively) injected with miR-467a-5p antagonist or control oligonucleotide.

**Figure 2.**
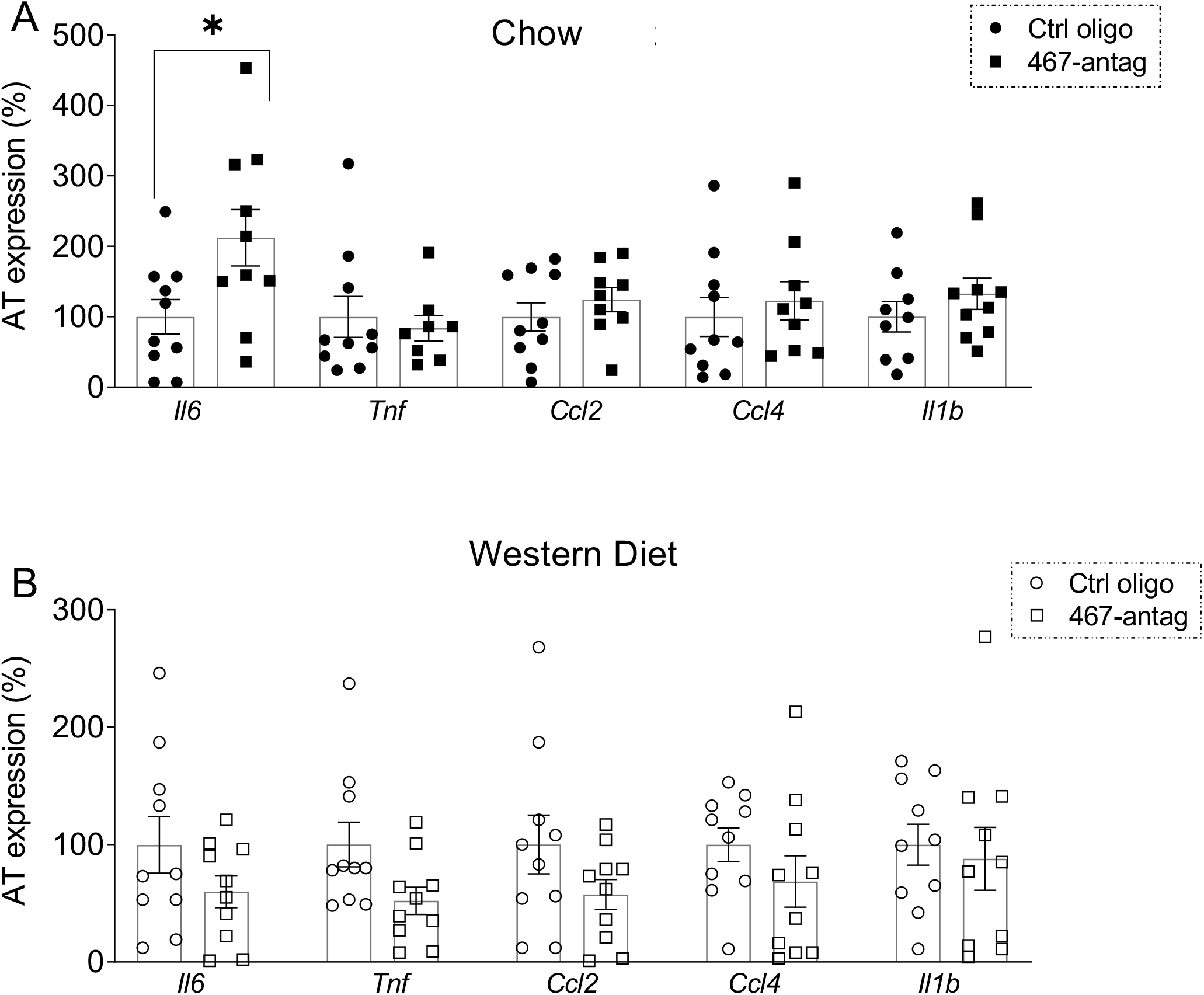
Inflammation in adipose tissue from WT mice on chow and Western diet. Effect of miR-467 antagonist on expression of pro-inflammatory markers (*Il6*, *Tnf*, *Ccl2*, *Ccl4*, *Il1b*) were assessed in WT chow (A) and Western diet (B) whole adipose tissue by RT-qPCR, normalized to β–actin. Data are relative to ctrl oligo. n=10 mice/group. \**P*<.05

Out of five cytokines, only *Il6* expression was statistically significantly increased by the antagonist in chow-fed mice (Figure 2A). Notably, all cytokine expression tended to be reduced in Western diet-fed mice in response to the antagonist (Figure 2B).

Western diet increased baseline expression of inflammatory markers in AT of WT mice injected with control oligonucleotide (Figure S4).

### 3.3 TSP-1 knockout does not prevent macrophages infiltration in mice injected with the miR-467a-5p antagonist

We reported that TSP-1 is a direct target of miR-467 ^21^ and the main mediator of miR-467 effects on cancer angiogenesis ^22^. *Thbs1^−/−^* mice were used, as described above, to determine whether macrophage infiltration is regulated by miR-467 through TSP-1 silencing. The miR-467a-5p antagonist did not prevent accumulation of AT macrophages from mice on chow or Western diet (Figures 3A, B). Macrophage infiltration in pancreas of *Thbs1^−/−^* mice was similar to infiltration of macrophages in pancreas of WT mice (Figures 3C, D).

**Figure 3.**
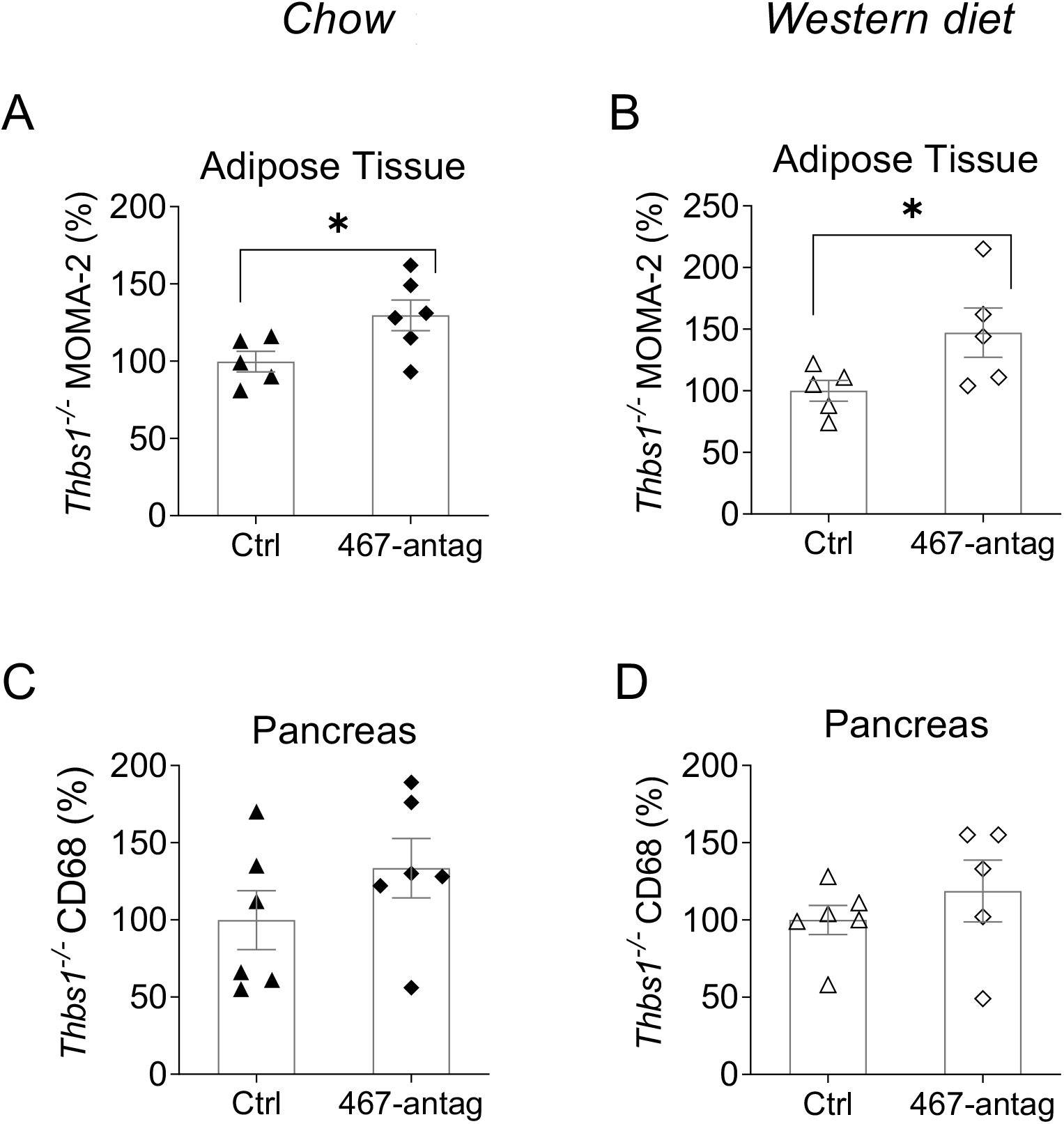
Macrophage accumulation in adipose tissue and pancreas from *Thbs1^−/−^* mice on chow or Western diet. Macrophage accumulation in adipose tissue from *Thbs1^−/−^* mice on chow diet (A) and Western diet (B) was determined by anti-MOMA-2 staining. Positive staining was normalized to mean adipocyte area for adipose tissue since adipocyte sizes were changed between groups. Macrophage accumulation in pancreas from *Thbs1^−/−^* mice on chow diet (C) and Western diet (D) was determined by anti-CD68 staining. Data are relative to ctrl oligo. n=7 mice/group. \**P*<.05

In *Thbs1^−/−^* mice, baseline AT macrophage infiltration was not affected by Western diet (Figure S5A), which was different from the effect of the Western diet in WT mice (Figure S2). Pancreas macrophage infiltration in *Thbs1^−/−^* mice on Western diet significantly increased ∼ 63.7% (Figure S5B, *P*=.03).

Similar to WT mice, *Thbs1^−/−^* blood monocyte numbers were not increased by the antagonist. The Western diet increased numbers of circulating monocytes and white blood cells (Figures S3D – F).

### 3.4 miR-467a-5p antagonist has no effect on macrophage infiltration and levels of inflammatory markers in liver

Hepatic inflammation and macrophage infiltration in liver are associated with development of hyperglycemia and IR ^38–40^. The levels of miR-467 were upregulated in liver of mice on Western diet (Figure S6A). We evaluated the expression of inflammatory and macrophage markers in liver (Figure S6). Expression of *Tnf* and *Il1b* were unaffected by the antagonist (Figure S6B, C and S6D, E) and macrophage infiltration was unchanged by the antagonist as shown by the expression of macrophage marker *Cd68* (Figure S6F, G).

### 3.5 Systemic injections of miR-467a-5p antagonist increase fasting insulin levels and decrease insulin sensitivity in chow-fed WT mice

In chow-fed mice, fasting blood glucose and insulin levels were measured at the end of the experiment. Inhibiting miR-467, using systemic injections of the antagonist, had no effect on fasting blood glucose levels in chow-fed mice but significantly increased fasting insulin levels (0.53 vs 1.01 mg/dL) (Figures 4A, B).

**Figure 4.**
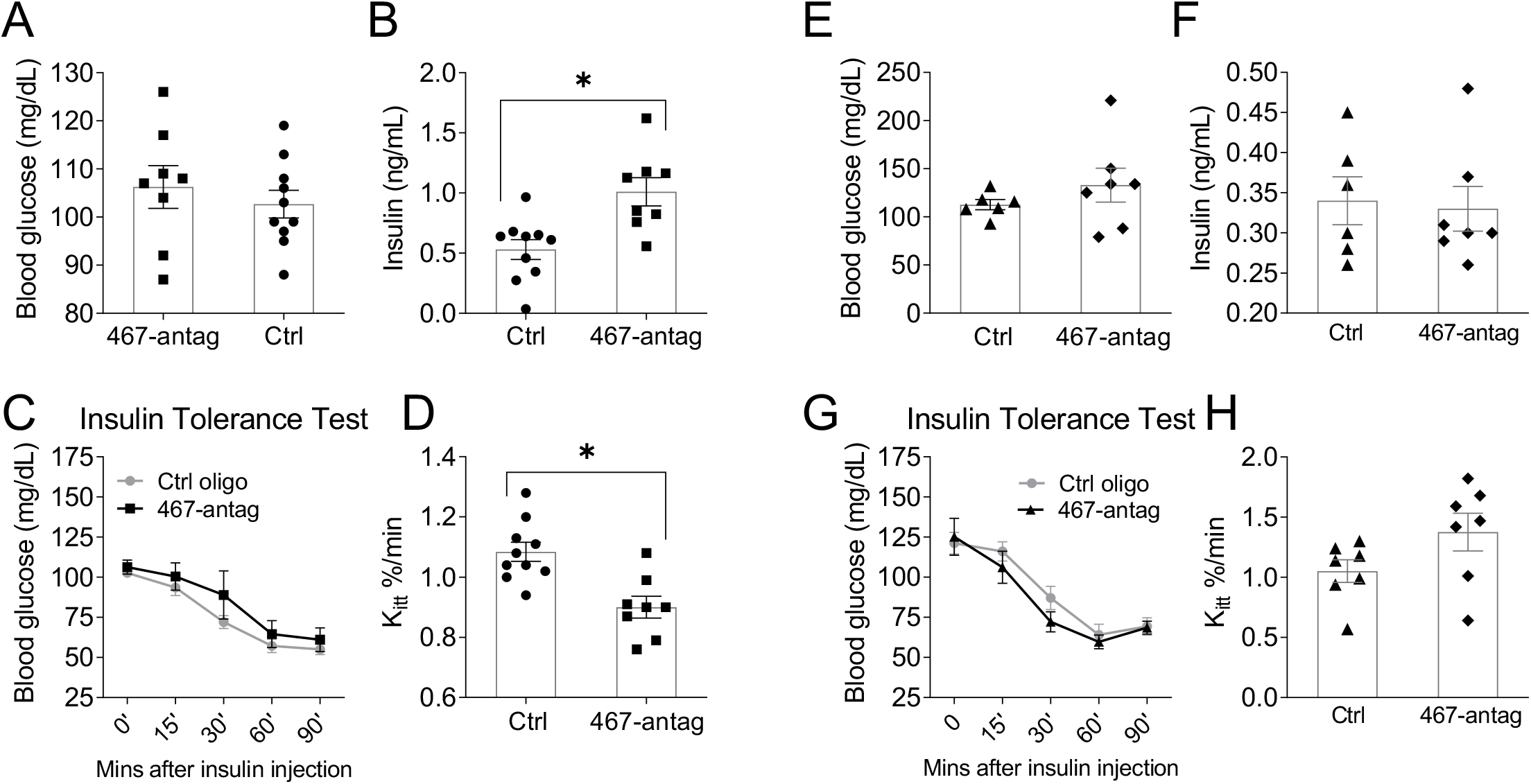
Inhibition of miR-467 increased fasting insulin and increased insulin resistance WT chow-fed mice, but not *Thbs1^−/−^* mice. Male WT (A – D) or *Thbs1*^−/−^ (E – H) mice began a chow diet at 4 weeks of age for 32 weeks. Starting at 5 weeks of age, mice received weekly injections of a control oligo or a 467-antagonist. Data are from end point. (A, E) Fasting blood glucose levels were measured with a glucometer. (B, F) Fasting insulin levels were measured by ELISA. (C, F) Time course for the intraperitoneal insulin tolerance test (ITT) were performed. (D, H) Rate constant for plasma glucose disappearance, K_itt_, from 0 – 60 minutes. \**P*<.05

In glucose tolerance tests (GTT), no changes were observed (Figure S7A). However, elevated glucose levels in antagonist-injected mice were observed during the ITT for all time points (Figure 4C).

Analyzing the rate of glucose disappearance from plasma (based on the K_itt_ analysis at 0 – 60 minutes, when the decrease in glucose levels was linear) revealed a significant decrease in glucose clearance in response to the antagonist (Figure 4D).

### 3.6 Systemic injections of miR-467a-5p antagonist do not affect mouse weight or blood lipid profile in chow-fed mice

Weight, HDL and LDL cholesterol, total cholesterol, and free cholesterol were measured. Antagonist injections had no effect on LDL (Figure S7D), but decreased HDL, total and free cholesterol, Figures S7C, E, F).

Similar to WT mice, weight was not affected by the antagonist in *Thbs1^−/−^* mice (Figure S7H). The levels of HDL, total and free cholesterol were unchanged in response to the antagonist (Figures S7I, K, L), but the LDL cholesterol levels were increased in mice injected with the antagonist in the absence of TSP-1 (Figure S7J).

### 3.7 TSP-1 knockout eliminates the effects of miR-467a-5p antagonist on insulin sensitivity in chow-fed mice

As with WT mice, fasting blood glucose and insulin levels were measured in *Thbs1*^−/−^ mice at the end of the experiment. *Thbs1^−/−^* mice were used in an identical experimental design as described above (Figure 4). Without TSP-1, there was no effect by the antagonist on blood insulin levels (Figure 4F). No differences were observed in GTT (Figure S7G). In *Thbs1^−/−^* mice, unlike in WT mice, blood glucose was not increased in the ITT in response to the antagonist at any time point (Figure 4G). Surprisingly, antagonist injections tended to improve IS, suggesting that other targets of miR-467 may become important in the absence of TSP-1. Loss of TSP-1 normalized, and even slightly increased, the plasma glucose disappearance rate in antagonist-injected mice (Figure 4H).

### 3.8 Systemic injections of miR-467a-5p antagonist increase blood glucose levels and decrease fasting insulin levels in WT mice on the Western diet

In Western-diet-fed mice, similar experiments revealed a different response to miR-467 inhibition and a loss of its protective function. As expected, mice on a Western diet developed diet-induced IR: fasting blood glucose levels were increased compared to chow-fed mice (102.7 ± 2.88 vs 117.9 ± 2.42, *P*<.001 Figure 5A vs 4A); insulin levels were twice higher in mice on Western diet (0.5301 ± .0820 vs .9327 ±. 0660, *P*=.0015 Figure 5B vs 4B). Antagonist injections further increased blood glucose levels by 20.85% (142.4 vs 117.9 mg/dL) in mice on the Western diet (Figure 5A) but significantly decreased blood insulin levels by 25.81% (0.69 vs 0.93 ng/mL, Figure 5B). Glucose clearance was delayed in the GTT (Figure 5C). The antagonist did not increase glucose levels in ITT, but a higher baseline blood glucose levels was detected compared to mice on chow in Figure 4A (Figure 5D). Analysis of the rate of glucose disappearance from plasma (K_itt_ analysis) revealed a significant increase in glucose clearance in antagonist-injected mice, which was opposite in chow-fed mice (Figure 5E vs 4D). These data suggest that, in mice on Western diet, the protective effect of miR-467 (decreased insulin levels and accelerated clearance of glucose from blood) is lost and even further counteracted by new, pro-IR effects of miR-467.

**Figure 5.**
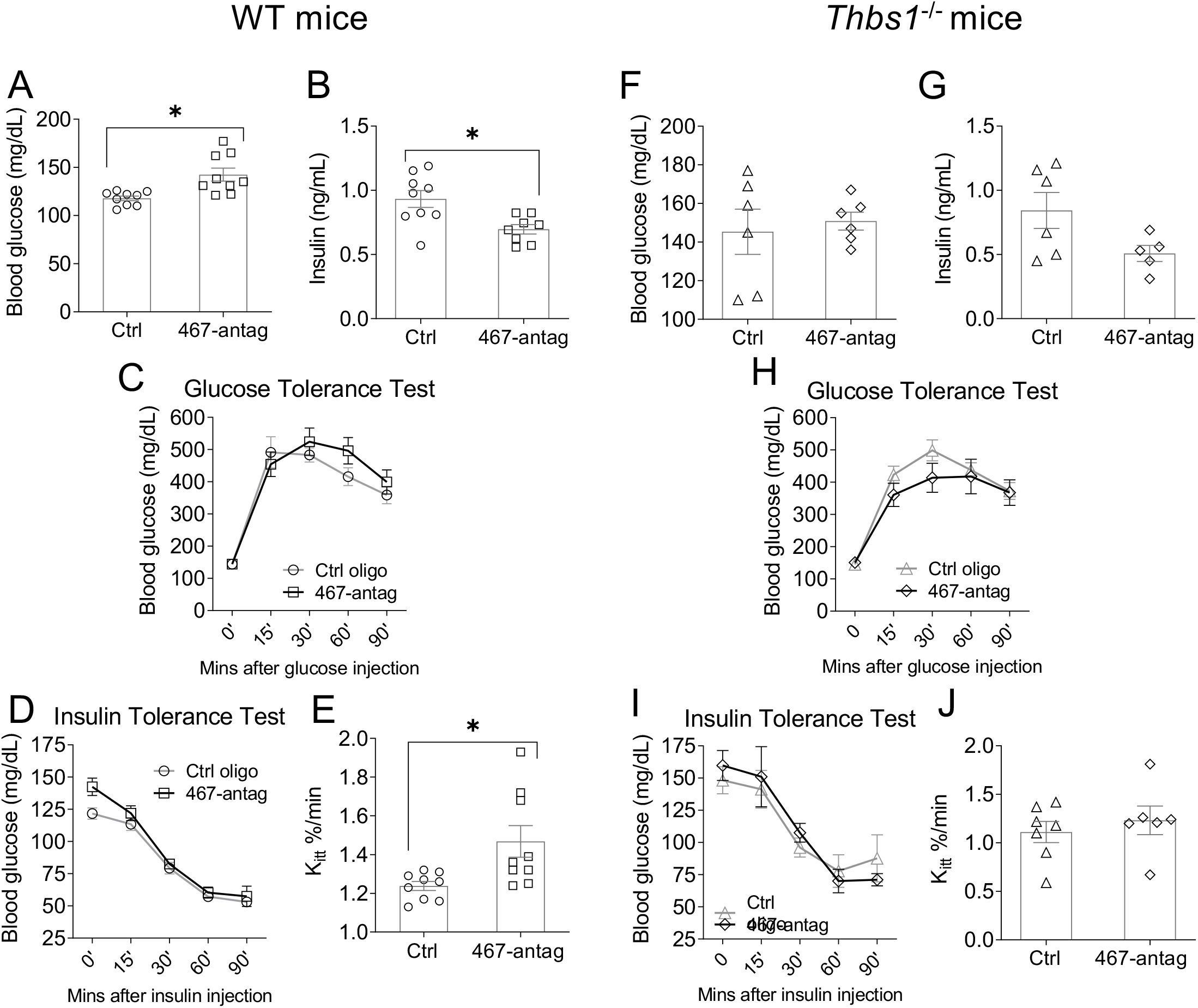
Inhibition of miR-467 in WT mice on a Western diet increased sensitivity to insulin, glucose clearance, and fasting blood glucose despite decreased fasting insulin. Male WT (A – E) or *Thbs1*^−/−^ (F – J) mice began a Western diet at 4 weeks of age for 32 weeks in an identical experiment as the chow-fed mice. Data are from end point. (A, F) Fasting blood glucose levels were measured with a glucometer. (B, G) Fasting insulin levels were measured by ELISA. Time course for the intraperitoneal glucose tolerance test (GTT) (C, H) and insulin tolerance test (ITT) (D, I) were performed. (E, J) Rate constant for plasma glucose disappearance, K_itt_, from 0 – 60 minutes. \**P*<.05

### 3.9 Systemic injections of miR-467a-5p antagonist do not affect mouse weight or blood lipid profile in WT mice on Western Diet

Similar to mice on the chow diet, the antagonist did not affect the weight or the lipoprotein cholesterol levels in WT mice on Western diet (Figures S8A – E). As was expected, the baseline weight and levels of HDL and LDL cholesterol, total and free cholesterol were increased by the diet itself (Figures S7C – F vs S8B – E).

### 3.10 The antagonist effect on insulin and glucose levels is lost in *Thbs1^−/−^* mice on Western

Similar to the effects in chow-fed mice, loss of TSP-1 in *Thbs1^−/−^* mice on Western diet abolished the antagonist effects on glucose and insulin levels and blood glucose clearance (Figures 5F – J). This indicates these functions of miR-467, and the effects of the antagonist depend on TSP-1 regulation. Loss of TSP-1 prevented increases in fasting blood glucose levels and decreases in fasting blood insulin levels (Figures 5F, 5G) that were observed in WT mice on Western diet Figures 5A, 5B). Additionally, loss of TSP-1 eliminated antagonist effects on blood glucose levels in GTT and ITT and plasma glucose clearance rate (Figures 5H – J).

### 3.11 Systemic injections of miR-467a-5p antagonist do not affect mouse weight or blood lipid profile in *Thbs1^−/−^* mice on Western Diet

Similar to WT mice, *Thbs1^−/−^* mouse weight, HDL and LDL cholesterol, and free and total cholesterol levels were not affected by the miR-467a-5p antagonist (Figures S8F – 8J). Weight and cholesterol levels were increased by the Western diet in both WT and *Thbs1^−/−^* mice (Figures S7 vs S8).

### 3.12. Effects of the miR-467 antagonist on the expression of glucose transporters (GLUT1, GLU2, GLUT4)

Expression of *Slc2a1*, a ubiquitous insulin-independent glucose transporter GLUT1, was measured in mouse AT, pancreas, and liver in mice on chow and Western diet injected with the antagonist or control oligonucleotide (Figures 6A – D).

**Figure 6.**
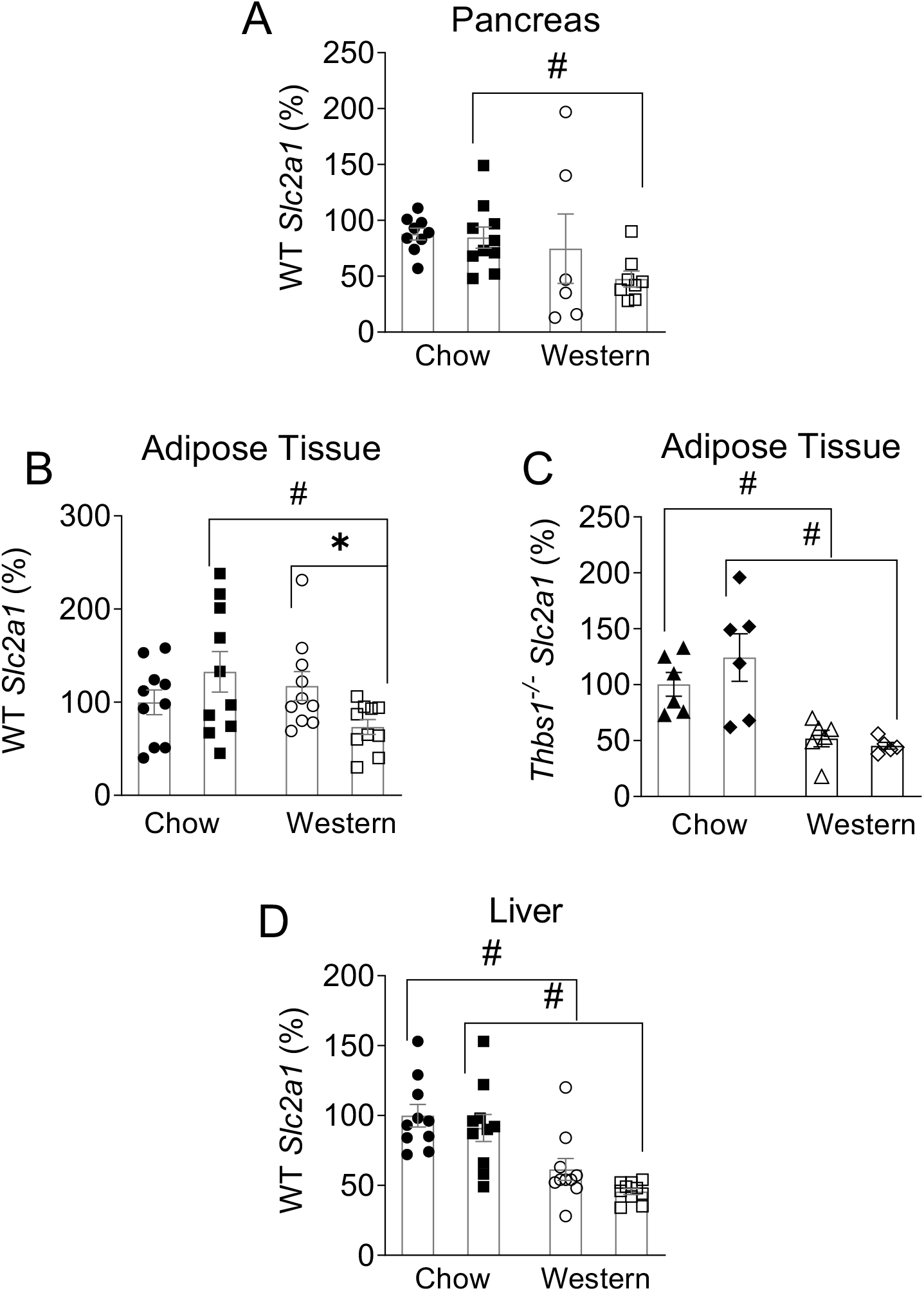
Effects of the miR-467 antagonist on the expression of glucose transporter *Slc2a1* (GLUT1). Expression of *Slc2a1*, a ubiquitous insulin-independent glucose transporter, GLUT1, was measured in WT pancreas (A), adipose tissue from WT (B) and *Thbs1*^−/−^ (C), and liver (D) in mice on chow or Western diet injected with the antagonist or control oligonucleotide. Data are relative to Chow ctrl. \**P*<.05 vs ctrl oligo, # *P*<.05 vs chow diet

No change in pancreas *Slc2a1* expression was detected in response to the miR-467 antagonist (Figure 6A).

*Slc2a1* expression in AT from Western-fed mice was decreased by the antagonist (Figure 6B), but the effect was lost in *Thbs1^−/−^* mice (Figure 6C), suggesting TSP-1 as a target in regulation of GLUT1.

In liver, *Slc2a1* expression was unaffected by the antagonist, but was decreased by the Western diet (Figure 6D).

In both pancreas and AT, there seemed to be a cumulative effect of the antagonist and Western diet: the expression was significantly decreased in antagonist-injected mice on Western diet compared to antagonist-injected mice on chow, without decreased expression in control oligonucleotide-injected mice (Figures 6A, B).

Expression of the major glucose transporters was also measured: *Slc2a2* (GLUT2) in pancreas and liver and *Slc2a4* (GLUT4) in AT (Figure S9). No changes were detected in these transporters in response to the antagonist. Western diet affected the expression in an organ- and transporter-specific manner: in pancreas, *Slc2a1* expression was lower in mice on Western diet (Figure 6A), while *Slc2a2* expression was increased (Figure S9A). In AT, *Slc2a1* was decreased Western diet-fed mice (Figure 6B), which was even more pronounced in *Thbs1^−/−^* mice (Figure 6C). AT *Slc2a4* expression was unchanged in either genotype, although in *Thbs1^−/−^* mice, expression tended to be lower in mice on the Western diet (Figures S9C, D).

### 3.13 miR-467a-5p in adipose tissue and the effects of the antagonist injections

To determine additional changes induced in AT by the antagonist, we evaluated the levels of miR-467 expression and TSP-1 protein, size of the adipocytes, and quantified ECM proteins (Figure S10). Differentiated 3T3-L1 (adipocyte-like cells) responded to high glucose (HG, 30 mM D-glucose) stimulation by increasing levels of miR-467a-5p by 21.8%±13.81 (Figure S10A, B; *P* = 0.02). However, *in vivo data,* AT miR-467 expression was unchanged in mice on Western diet (Figure S10C) at the end of the experiment, possibly reflecting the transient upregulation of miR-467 in response to hyperglycemia.

To assess how TSP-1 protein levels were changed in AT, sections were stained with an anti-TSP-1 antibody (Figure S10D). Area stained with anti-TSP-1 was decreased in Western diet-fed mice by 71.40% in control group and 49.5% in antagonist-injected mice (Figure S10D), apparently reflecting the increase in adipocyte size (Figure S10E, F and reduction in overall fraction of area between AT cells. As expected, the antagonist tended to rescue TSP-1 levels in WT mice by 27.30% on chow and 48.97% on Western diet.

Hypertrophic adipocytes contribute to the release of inflammatory cytokines, immune cell recruitment, and impaired insulin sensitivity. AT sections were H&E stained to quantify cell sizes (Figures S10E, F). 4864 to 6749 adipocytes per animal were analyzed. Mean areas and perimeters of adipocytes were increased by the Western diet and tended to increase in response to miR-467a-5p antagonist injections (Figures S10E, F).

Fibrosis and ECM deposits between cells in AT affect remodeling, growth, and function of adipocytes ^32,39,41^. To evaluate changes of ECM amounts in AT, sections were stained with Masson’s Trichrome to assess ECM levels. There was no difference in staining between the mouse groups (Figure S10G, H).

### 3.14 miR-467a-5p in pancreas and the effects of the antagonist injections

To evaluate other effects of the antagonist in the pancreas, we examined miR-467 and TSP-1 levels, islet area, and vascularization (Figure S11).

Mouse pancreatic islets β-cells (βTC-6) were stimulated with high glucose (HG) and miR-467a-5p levels were measured. Glucose-stimulated cells significantly increased miR-467a-5p expression by 27.7 ± 4.93% (Figure S11A, *P* = .03). The antagonist significantly decreased βTC6 expression of miR-467 by 81% (Figure S11B, *P* = .002). In the *in vivo* experiment, the mean value of pancreatic miR-467a-5p was increased two-fold on the Western diet (208.1% ± 173.9 vs. 108.6% ± 39.04 in chow diet), but was not statistically significant (Figure S11C).

Sections of pancreas were stained with the anti-insulin antibody and counterstained with hematoxylin (Figure S11D). The total islet area in the pancreas was unchanged in any of the mouse groups.

We have previously reported that miR-467a-5p promotes angiogenesis as a result of regulation of production of its target, thrombospondin-1 (TSP1) ^21,22^. Thus, we assessed the potential effect of miR-467a-5p antagonist on vascularization and TSP-1 in the pancreas by immunohistochemistry with anti-vWF, anti-α-actin, and anti-TSP-1 antibodies. There was no change in the vascular cell markers, vWF (marker of endothelial cells) or α-actin (marker of vascular smooth muscle cells) in miR-467a-5p antagonist-injected mice or in response to the Western diet (Figures S11E, F). We also evaluated TSP-1, a target of miR-467, in the pancreas. At the end of the experiment, TSP-1 levels were not affected by miR-467 antagonist injections or by the diet (Figure S11G).

### 3.15. Expression of gluconeogenesis gene expression in liver of mice injected with miR-467 antagonist

To evaluate whether changes in blood glucose were mediated due to regulation of gluconeogenesis, we examined expression of key gluconeogenesis enzymes in liver (Figure S12). *G6pc* encodes glucose-6-phosphatase, a regulator of conversion of glucose 6-phosphate to glucose. Liver *G6pc* expression was decreased by the antagonist (significant in mice on the Western diet) (Figure S12A). Thus, *G6pc* could not be responsible for the higher levels and slower clearance of blood glucose from Figure 4. *G6pc* expression was significantly decreased by the Western diet. *Fbp1* is an enzyme catalyzing the hydrolysis of fructose 1,6-bisphosphate to fructose 6-phosphate and acting as a rate-limiting enzyme in gluconeogenesis. Liver *Fbp1* was unaffected by antagonist injections but, like *G6pc*, was decreased by the Western diet (Figure S12B).

### 3.16 High glucose upregulates miR-467a-5p in macrophages

Murine macrophages (RAW 264.7) and differentiated human monocyte (THP-1) cell lines were stimulated with high glucose (HG, 30mM D-glucose and 30mM L-glucose), and miR-467a-5p expression was measured. miR-467a-5p was upregulated by 20.6% ± 14.31 in RAW 264.7 and by 540.3% ± 559.5 in THP-1 in response to HG concentrations (Figures 7A,C, respectively). Increased miR-467a-5p levels were associated with a decrease in TSP-1 production by the cells (Figures 7B, 7D, *P* < .05). Consistent with the mechanisms of miR-467a-5p upregulation by HG previously described by us ^21,31^, both D-glucose and the biologically inactive L-glucose had similar effects: HG upregulated miR-467a-5p expression and decreased secreted TSP-1, which indicated the osmolarity change as a stimulus for miR-467a-5p upregulation in macrophages.

**Figure 7.**
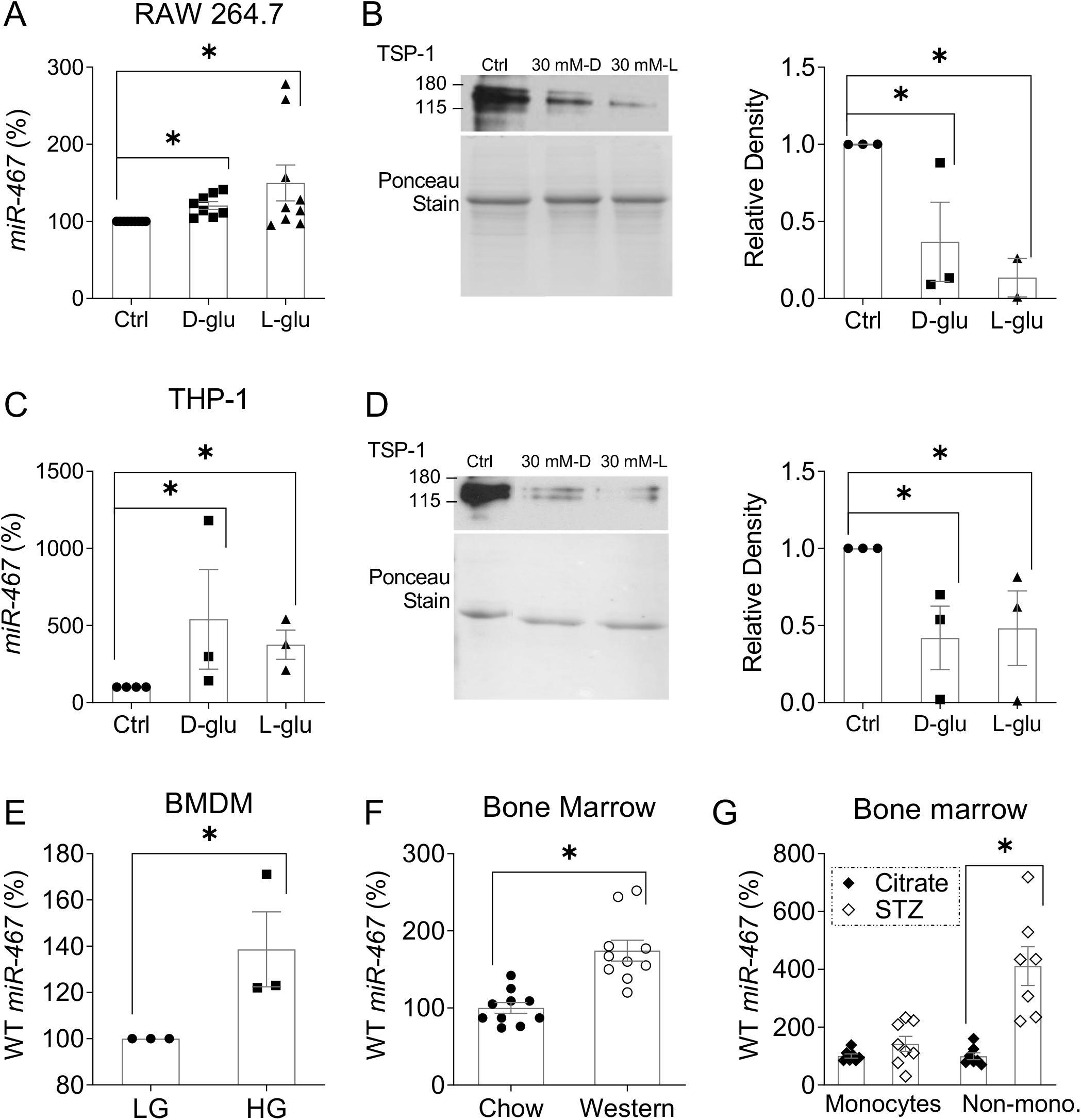
High glucose upregulates miR-467 in macrophages. (A, C) Expression of miR-467 after 3 hours of glucose stimulation in cultured mouse macrophages (RAW 264.7) and differentiated human monocyte (THP-1) cell lines was measured by RT-qPCR and normalized to β-actin, n=3-8 independent replicates. (B, D) TSP-1 secretion was assessed in cell supernatants after 24 hrs of glucose stimulation by Western Blot. Quantification of densitometry is shown and relative to the control, n=3 independent replicates. \**P*<.05 vs ctrl. (E) miR-467 expression in cultured WT bone marrow-derived macrophages (BMDM) 6 hours post glucose stimulation. Data are relative to low glucose (LG) ctrl samples, n=3 independent replicates. \**P* <.05 (F) miR-467 expression in whole bone marrow (BM) from WT mice on chow or Western diet for 32 weeks, n=10 mice/group. Data are relative to chow mice \**P*<.05 (G) miR-467 expression in BM monocytes or non-monocytes from male BALB/c mice injected with STZ to induce diabetes, or a citrate buffer control. Data are relative to citrate buffer ctrl. n=10 mice/group. \**P*<.05 vs citrate buffer ctrl

In cultured mouse bone marrow-derived macrophages (BMDM), miR-467a-5p was significantly upregulated in response to glucose by 38.7% ± 28.01 (Figure 7E, *P* = .05). miR-467a-5p levels were increased in bone marrow (BM) by 74.13% in Western diet-fed mice (174.3% ± 42.72 vs. 100.1% ± 21.95 in chow, *P* < 0.001) compared to chow-fed mice (Figure 7F). miR-467a-5p levels were increased 4-fold in the non-monocytic fraction in bone marrow from STZ-treated hyperglycemic BALB/c mice and tended to be increased in the monocytic fraction (non-monocytes: 411.1% ± 176.9 in STZ vs. 99.86 ± 32.01 in citrate buffer, Figure 7G, *P* = 0.001).

### 3.17 miR-467a-5p mimic and antagonist regulate production of inflammatory signals by the cultured macrophages

Cultured BMDMs from WT mice were transfected with a miR-467a-5p mimic or 467-antagonist as described in Methods and expression of *Tnf*, *Il6*, *Ccl2*, and *Ccl4* were measured. Expression of all four cytokines was increased by HG (Figures S13A, B), but miR-467a-5p mimic had no additional effect (Figure S13A).

Inhibiting miR-467a-5p with the antagonist in BMDMs from WT mice prevented the upregulation of *Tnf* and *Ccl4* in response to HG (Figure S13B), suggesting that these two cytokines are regulated by HG through the miR-467-dependent mechanism, while others are not.

When cultured BMDMs isolated from *Thbs1^−/−^* mice were transfected with the miR-467a-5p mimic or antagonist, upregulation by high glucose was similar to BMDMs from WT mice (Figures S13C, D), except for *Ccl4* which was not upregulated by HG in the absence of TSP-1. The increase in *Tnf* by HG was still blunted by the antagonist, suggesting this effect is not dependent on TSP-1 and that miR-467 uses multiple targets in regulating inflammation. We did not observe a difference in basal levels of cytokines between WT and *Thbs1^−/−^* cells (not shown).

### 3.18 Differential effects of the miR-467a-5p antagonist on plasma levels of inflammatory cytokines

Plasma levels of MCP-1, IL-10, CXCL1, and VEGF-A were measured in WT and *Thbs1^−/−^* mice on chow or Western diet (Figures S13E – H). MCP-1, IL-10, and CXCL1 levels were significantly increased by the Western diet in both mouse genotypes (Figures 13E – G). The effects of the Western diet on the levels of cytokines were specific: VEGF-A was not increased (Figure S13H).

In WT mice, MCP-1 levels were increased by the miR-467a-5p antagonist on the Western diet (Figure S13E, *P* = 0.05) but not in *Thbs1^−/−^* mice. The antagonist tended to decrease the levels of IL-10 in WT mice on the Western diet (Figure S13F, *P* = 0.06), but this effect was lost in the *Thbs1^−/−^* mice. Thus, these two markers were regulated by miR-467 and TSP-1.

There was no effect of the antagonist or TSP-1 deletion on CXCL1 or VEGF-A levels (Figures S13G, H), suggesting that these two markers are not regulated by miR-467 or TSP-1.

### 4. DISCUSSION

The sequence and causality of events in development of IR are still poorly understood ^42^, and physiological mechanisms normally preventing development of IR are unclear. Dietary factors increasing blood glucose and insulin production may induce IR ^43–46^, and inflammation and infiltration of metabolically active tissues with macrophages are recognized as important and causative factors in IR development ^32–37^. Here we report miR-467 decreases blood insulin level and accelerates glucose clearance: the injections of miR-467 antagonist increased fasting insulin levels and reduced insulin sensitivity and glucose clearance from the blood. Inhibition of miR-467 in chow-fed mice raised insulin levels up to those of mice on the Western diet, and Western diet, rich in fats and sugars, results in a loss of this physiological function of miR-467, either due to the presence of other unidentified mRNA targets induced by the Western diet or due to other cellular mechanisms activated by the Western diet, which are counteracting miR-467 effects.

Metabolic disorders are often thought to be a direct consequence of the weight gain and changes in the lipoprotein profile ^47^. When we inhibited miR-467a-5p, the changes in the blood glucose and insulin levels were uncoupled from the weight gain and impairment of lipoprotein metabolism, suggesting that the effect of miR-467a-5p is not mediated by change in weight or cholesterol.

One of the potential reasons identified may be the decreased expression of an insulin-independent glucose transporter GLUT1, a main glucose transporter regulating insulin production in human pancreas. The decreased expression of *Slc2a1* in response to the antagonist injections in mice on Western diet coincided with lower blood insulin levels, higher glucose levels, and increased glucose clearance from plasma. Regulatory regions of *Slc2a1* (GLUT1) mRNA do not have predicted target sites for miR-467, thus, the regulation by miR-467 is most likely indirect. The effect on *Slc2a1* expression in the absence of an effect on other glucose transporters suggests that the regulation may not be associated with change in insulin sensitivity and that regulation of blood insulin levels by miR-467 may be secondary.

Inflammation and the expression of glucose transporters and key enzymes of gluconeogenesis were examined in livers, but no changes were detected in response to the antagonist injections, confirming that the effect of the antagonist is associated with the regulation of glucose clearance rather than glucose production.

We previously reported that TSP-1 transcript is a target of miR-467 and mediates miR-467 effect on angiogenesis. Interestingly, all effects of the antagonist on the blood glucose and insulin levels were lost in *Thbs1^−/−^* mice, suggesting that TSP-1 is the main target of miR-467, and the differential regulation in chow-fed and Western-diet-fed mice is downstream of TSP-1. TSP-1 is a known regulator of insulin sensitivity and metabolic disorder ^18–20,48,49^.

In adipose tissue, pro-inflammatory molecules are released by adipocytes and activated macrophages to promote insulin resistance ^50–53^. The inhibition of miR-467a-5p increased infiltration of macrophages in the adipose tissue and in the pancreas, suggesting that miR-467a-5p prevents inflammation. Additionally, our results and reports from others stress the importance of ECM, and TSP-1 (a target of miR-467a-5p) specifically, and other TSPs, in the recruitment of inflammatory cells into tissues ^18–20,54–58^. The increase in macrophage infiltration in adipose tissue was associated with the increased *Il6* levels, which was lost in *Thbs1*^−/−^ mice. However, the levels of *Tnf, Ccl2*, *Ccl4*, and *Il1b* were not changed by the antagonist injections.

The reduction of inflammation in the obese AT in response to the miR-467 antagonist may be due to a significant decrease in GLUT1 (*Slc2a1*) expression. Bone-marrow-derived macrophages isolated from mice with a myeloid-specific knockout of GLUT1 (*Slc2a1*) were “metabolically reprogrammed” such that they were unable to uptake glucose properly and had a decreased inflammatory phenotype ^59^. In tissues from Western diet-fed mice, there may be additional miR-467 targets not expressed in chow-fed mice; these may modify the antagonist effects, thus abolishing the inflammation and IR protection by miR-467.

The role of miR-467 is not limited to regulation of local inflammation in tissues: the effect of miR-467 inhibition on systemic inflammation was observed by changes in plasma levels of MCP-1 and IL-10. Both were increased in mice on Western diet, and MCP-1 was further increased upon inhibition of miR-467. In *Thbs1^−/−^* mice, the antagonist had no effect, suggesting that both cytokines in plasma are regulated by miR-467 through a TSP-1-dependent pathway.

miR-467a-5p was upregulated by high glucose in primary bone-marrow-derived macrophages (BMDMs), macrophage-like cell lines, and inflammatory blood cells from the bone marrow, suggesting that macrophages aid in regulating miR-467a-5p-dependent pathways, and macrophages infiltration may enhance the significance of the pathway in metabolically active tissues. Increased miR-467a-5p levels coincided with the inhibition of TSP-1 production by macrophages, as we observed previously in other cell types ^21,22,31^.

miR-467a-5p regulated the pro-inflammatory functions of cultured BMDMs: cytokine expression of *Tnf, Il6, Ccl2, and Ccl4* was upregulated by high glucose (HG). Only *Ccl4* and *Tnf*, were upregulated in cultured macrophages by HG through the miR-467-dependent mechanism; their upregulation was prevented by the miR-467 antagonist.

Only *Ccl4* appears to be regulated through TSP-1 pathway: upregulation, and the effect of miR-467 antagonist, were lost in BMDMs from *Thbs1^−/−^* mice. These results suggested that inflammation is regulated by miR-467a-5p through multiple targets and in a cell-specific manner in various cell types.

Our results unveil the physiological role of miR-467a-5p: when this miRNA is upregulated by high blood glucose ^21,22,31^, it protects against the development of IR and inflammation in response to high glucose. Interestingly, this protection is lost under a long-term Western diet, underscoring the negative effects of this chronic stressor.

## AUTHOR CONTRIBUTIONS

JG performed experiments, analyzed experimental data, developed experimental plan, and wrote the manuscript. IK performed experiments, analyzed experimental data, participated in discussion of the results and preparation of the manuscript. RY performed immunohistochemistry experiments, analyzed experimental data, and participated in discussion of the results and the plan for the manuscript. DV performed animal experiments, contributed to the discussion of the results and preparation of the manuscript. AV analyzed the adipocyte size, developed the program and the plan of for these analyses, and participated in the discussion of the result and the manuscript. LS performed experiments in cultured cells, analyzed experimental data, and participated in discussions of the results and of the manuscript. OS-A sponsored the project, developed the experimental design, participated in generation of immunohistochemistry data, analyzed experimental data, and prepared the manuscript.

The first author, JG and the corresponding author, OS-A, take full responsibility for the work as a whole, including (if applicable) the study design, access to data, and the decision to submit and publish the manuscript.

## ACKNOWLEDGEMENTS

This work was supported by R01CA177771 and R01HL117216, to OS-A and 17PRE33660475 from the American Heart Association (PI: JG, Sponsor: OS-A).

## DATA AVAILABILITY STATEMENT

The data that supports the findings of this study are available from the corresponding author upon reasonable request.

## 2. MATERIALS AND METHODS

### 2.1 Experimental animals

Animal procedures were approved by the Institutional Animal Care and Use Committee. Up to 5 mice were housed per cage and allowed access to food *ad libitum.* Male WT C57BL6 (n=10/group) or *Thbs1^−/−^* (n=7/group) mice were fed a chow or Western diet (TD.88137, 40-45% kcal from fat, 34% sucrose by weight, Envigo) starting at 4 weeks of age and injected weekly with a miR-467a-5p antagonist (2.5 mg/kg body weight) (or a control oligonucleotide that does not have predicted targets in the mouse and human genomes^22,28^), intraperitoneally, starting at 5 weeks of age until the end of the experiment. Body weight was measured weekly.

### 2.2 miR-467a-5p mimic and the miR-467a-5p antagonist

The miR-467a-5p mimic and the control oligonucleotide were purchased from Dharmacon. Cholesterol conjugated miR-467a-5p was modified by tagging a fluorophore (DY547) and a cholesterol moiety. The custom LNA-modified miR-467a-5p antagonist (TacaTGcaGGcacTTa) and a control oligonucleotide (TTTaGaccgaGcgTGt) were from Qiagen.

### 2.3 Glucose and insulin tolerance tests (GTT and ITT)

GTT and ITT were administered after overnight fasting. Glucose (2 g/kg body weight) or insulin (50 µg/kg) (Sigma) were injected intraperitoneally. Blood glucose levels were measured 0 – 180 min after injections using an AlphaTRAK glucometer. The glucose removal rate (Kitt), expressed as % / minute, was calculated using the following formula: (0.0693/(t1/2) x 100. Plasma glucose (t1/2) was calculated from the slope of the least squares curve analysis during the period when plasma glucose concentrations decreased linearly, from 0 – 60 min ^29,30^.

### 2.6 Immunohistochemical staining

Visceral (omental) adipose tissue and pancreas were fixed in 4% formaldehyde (Electron Microscopy Sciences) for 24 hours, transferred into 70% ethyl alcohol, and embedded in paraffin blocks. Two 5 μM sections of tissue per animal were stained with Hematoxylin (Ricca), Eosin (Protocol), Masson’s trichrome, or specific antibodies. H&E stained sections of adipose tissue were analyzed by ERT Imaging (Cleveland, OH) to determine adipocyte sizes.

Using VECTASTAIN ABC-HRP Kit (Vector Labs), sections were stained with anti-CD68 (biotinylated clone FA-11, 1:10, AbD Serotec), anti-Insulin (1:100, Dako), MOMA-2 (1:25 AbD Serotec), anti-vWF (1:400, Dako), anti-α-actin (clone ab5694 1:200, Abcam), or anti-TSP-1 Ab4 (clone 6.1 1:100, Thermo). Secondary antibodies were included in the species-specific kit and were followed by ImmPACT DAB peroxidase substrate (Vector Labs). Slides were scanned using Leica SCN400 or Aperio AT2 at 20X magnification. Quantification of positive staining was performed using Photoshop CS2 (Adobe) or Image Pro Plus (7.0).

### 2.7 Cell culture

RAW264.7, THP-1, βTC6 and 3T3-L1 cells were purchased from ATCC and cultured according to ATCC directions. THP-1 cells were differentiated in 100 nM PMA (Sigma) for 3 days before glucose stimulation. 3T3-L1 cells were differentiated at 80% confluency with 1µM Dexamethasone, 0.5 mM IBMX, and 1 µg/mL Insulin (all from Sigma).

### 2.8 Isolation of bone marrow-derived macrophages (BMDM)

Bone marrow was collected from femurs and tibia as described in ^31^. Macrophages were differentiated from whole bone marrow using 30 ng/mL MCSF (Biolegend) for 4 days, followed by 15 ng/mL MCSF for 3 days.

### 2.9 Glucose stimulation of RAW264.7, differentiated THP-1, βTC6, and BMDM

Up to 1.0 x10^6^ cells were plated in complete media in 6-well plates (Corning). Once glucose levels reached the fasting level (90 mg/dL) as measured using AlphaTRAK glucometer, cells were stimulated with 30 mM D-glucose High Glucose, “HG” (Sigma) for 6 hours (RAW 264.7 and BMDM), 3 hours (3T3-L1) or 30 minutes (βTC6).

### 2.10 Transfection of cultured cells

Transfection of the miR-467a-5p antagonist and its control oligo were aided with Oligofectamine (Invitrogen) for 24 hours. Successful transfection with the cholesterol-modified miR-467a-5p mimic was confirmed by fluorescence 24 hours post-transfection using an inverted microscope DMI6000SD (Leica).

### 2.11 Oil Red O Staining

Differentiated 3T3-L1 cells were washed with 1X PBS and fixed in 10% formalin (Electron Microscopy Sciences) for 15’ at room temperature (RT), washed with 60% isopropanol (Sigma), and stained in the Oil Red O solution for 10’ at RT.

### 2.12 RNA Extraction and RT-qPCR

RNA was isolated using Trizol reagent (Thermo). Organs were flash frozen in liquid nitrogen and homogenized in Trizol. RNA was quantified using Nanodrop 2000 (Thermo).

To measure miR-467a-5p expression, 1 – 2.5 μg of total RNA was first polyadenylated using NCode miRNA First-Strand cDNA Synthesis kit (Invitrogen) or miRNA 1st strand cDNA synthesis kit (Agilent). Real-time qPCR amplification was performed using SYBR GreenER™ qPCR SuperMix Universal (Thermo) or miRNA QPCR Master Mix (Agilent). The miR-467a-5p primer (GTA AGT GCC TAT GTA TATG) was purchased from IDT.

To measure expression of inflammatory markers, 1 – 2 μg of total RNA was used to synthesize cDNA using the SuperScript First-Strand cDNA Synthesis System for RT-PCR (Invitrogen). Real-time qPCR was performed using TaqMan primers for *Tnf*, *Il6*, *Ccl2*, *Il1b*, *Il10*, *Ccl4, Cd68, Slc2a1, Slc2a2, Slc2a4, G6pc, Fbp1* (Thermo) and TaqMan Fast Advanced Master Mix (Thermo). Ct values were determined as described previously ^32^.

### 2.13 Statistical analysis

Data are expressed as the mean value ± Se (standard error). Statistical analysis was performed using GraphPad Prism 5 Software. Student’s t-test and ANOVA were used to determine the significance of parametric data, and Mann-Whitney test was used for nonparametric data. A *P*-value of <.05 was considered statistically significant.

## Supplementary Figure Legends

**Figure S1.**
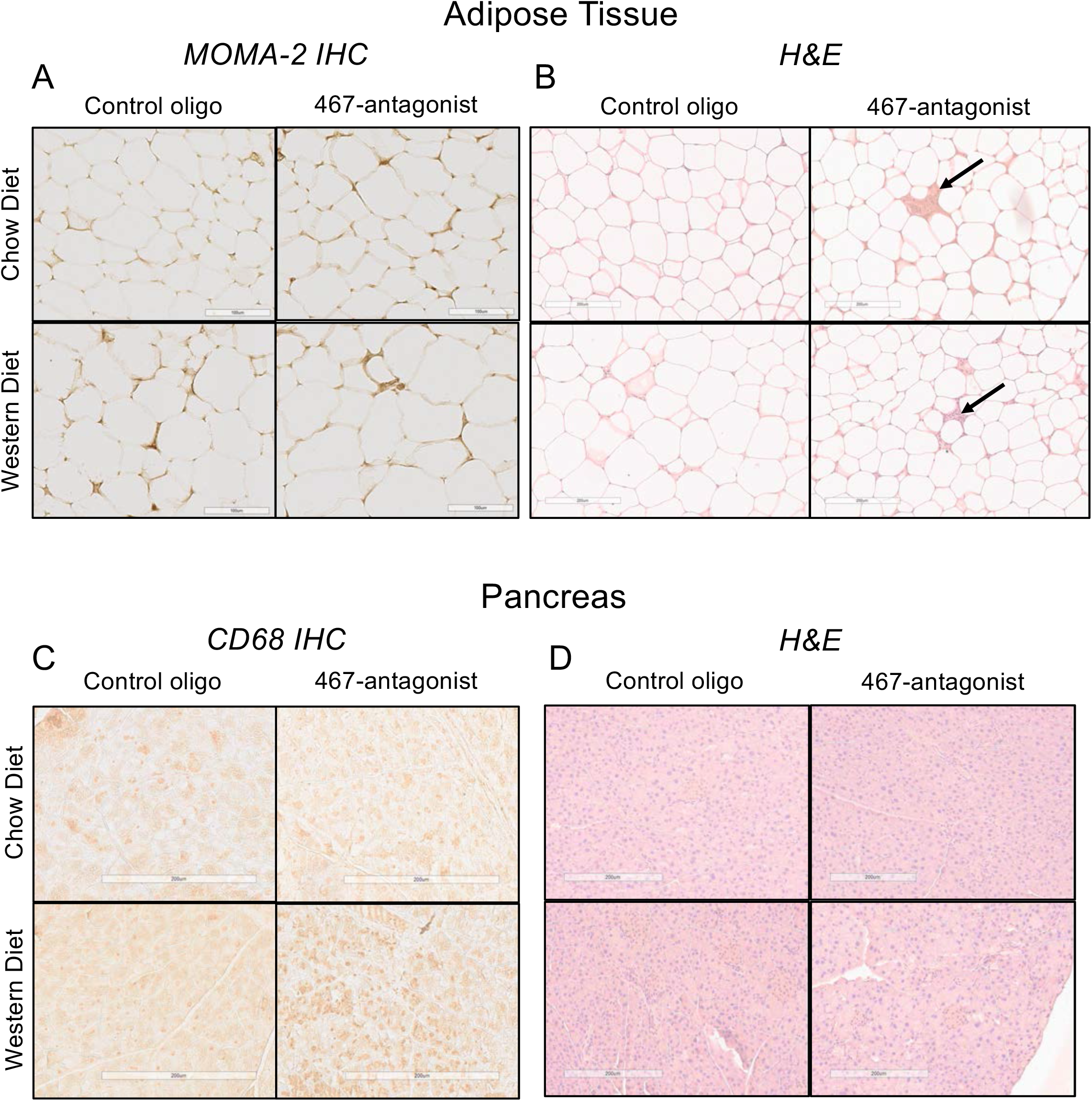
Representative images of macrophage accumulation markers and tissue structure in WT adipose tissue and pancreas. Representative images in WT adipose tissue (A, B) or pancreas (C, D) are shown. Adipose tissue was stained with anti-MOMA-2 antibody (A) or H&E (B). Pancreas was stained with anti-CD68 antibody (C) or H&E (D). Scale bars at 200µM (MOMA-2 IHC adipose tissue scale bars at 100 µM). n=10 mice/group. Arrows show the crown structures.

**Figure S2.**
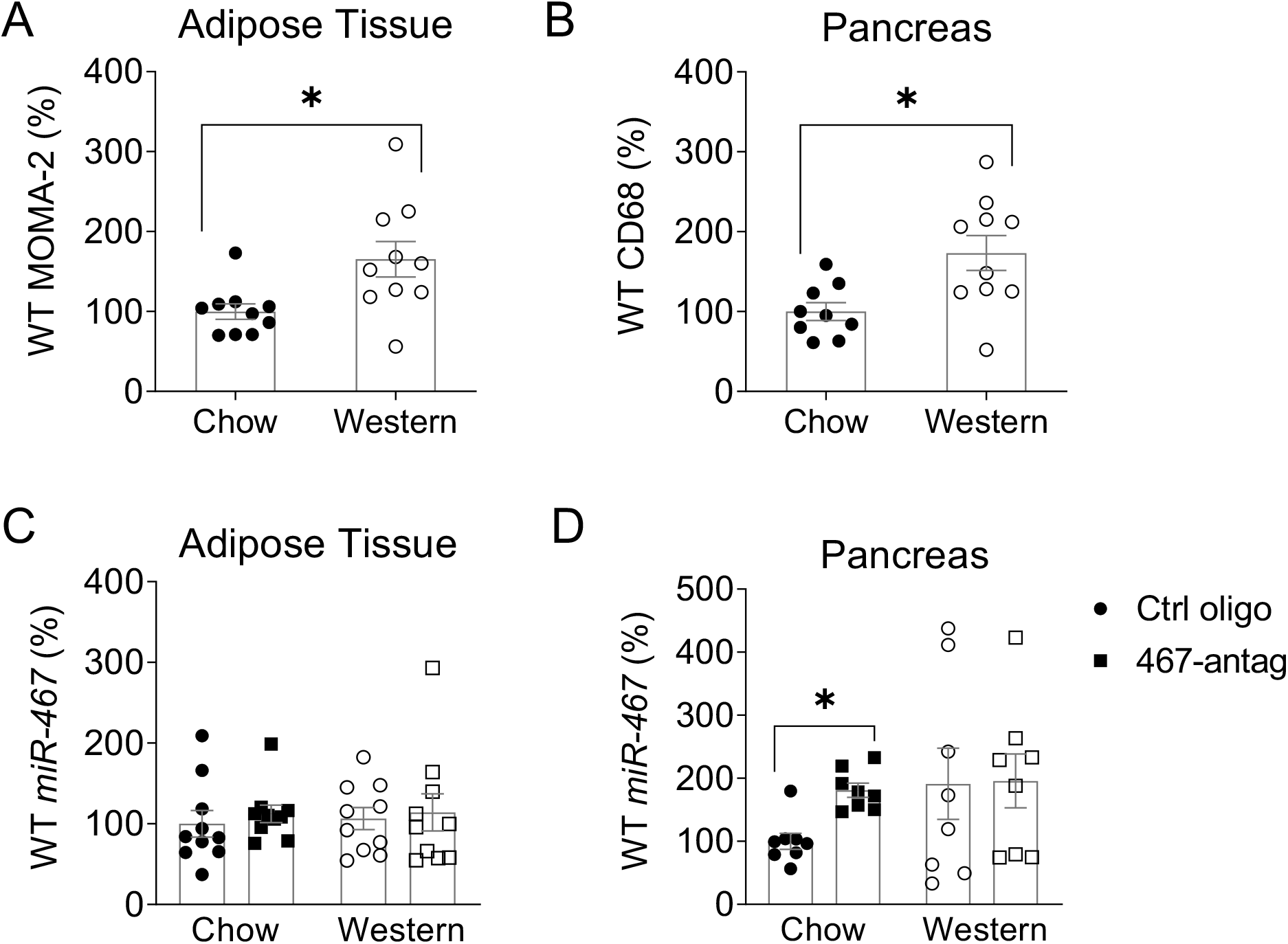
Effect of Western diet on macrophage accumulation in adipose tissue and pancreas in WT mice. Effects of the Western diet on macrophage accumulation in adipose tissue and pancreas was determined by (A) anti-MOMA-2 or (B) anti-CD68 in WT mice (injected with the control oligonucleotide), respectively. In AT, positive staining was normalized to mean adipocyte area for adipose tissue since adipocyte sizes were changed between groups. n=10 mice/group. Data are normalized to Chow ctrl diet average staining. \**P*<.05

**Figure S3.**
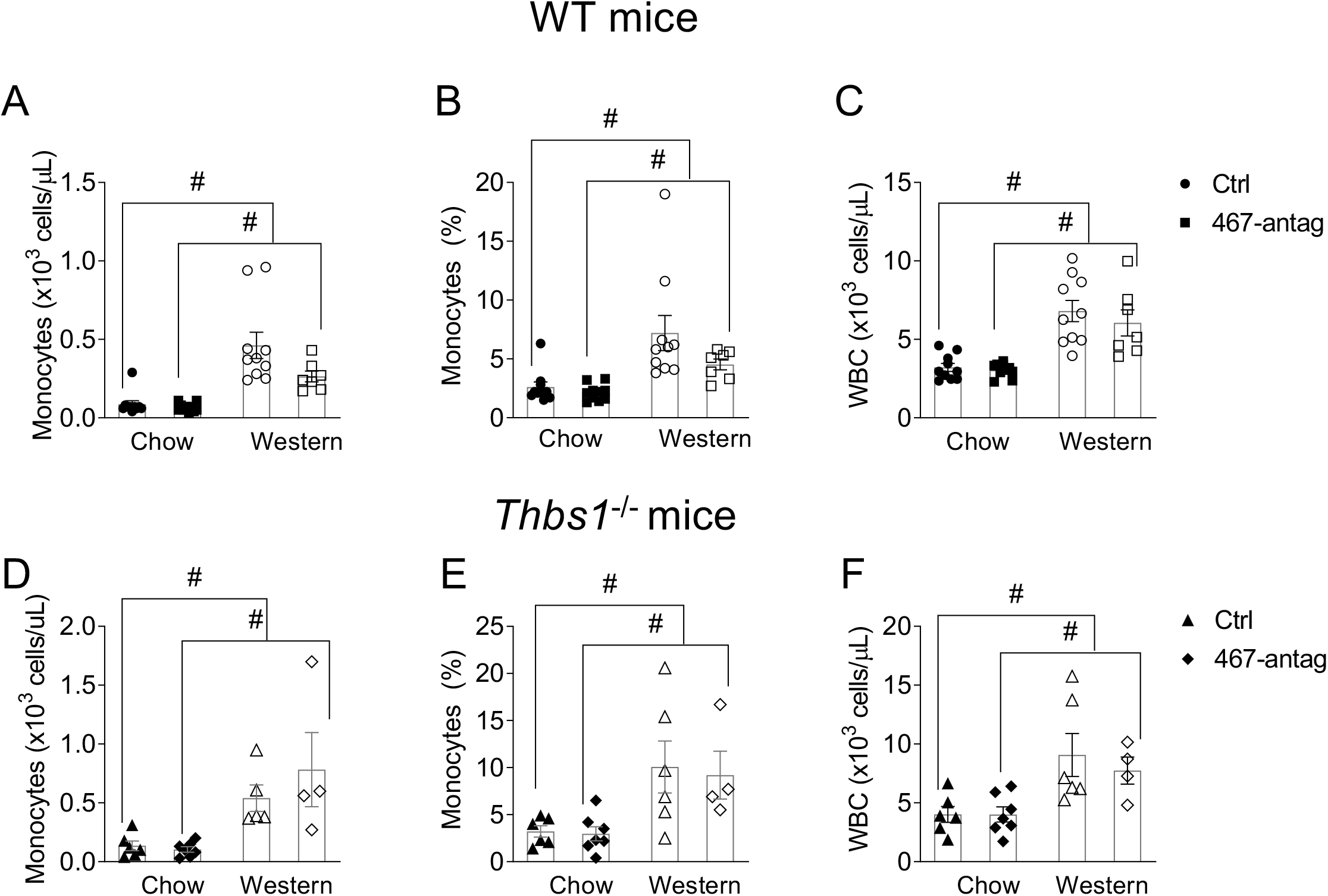
miR-467 antagonist has no effect on circulating monocytes or WBC in WT or *Thbs1*^−/−^ mice. Whole blood was collected at end point and analyzed on a hematology analyzer to determine circulating numbers of monocytes and WBCs in WT (A – C) or *Thbs1^−/−^* mice (D – E). # *P*< .05 vs chow diet.

**Figure S4.**
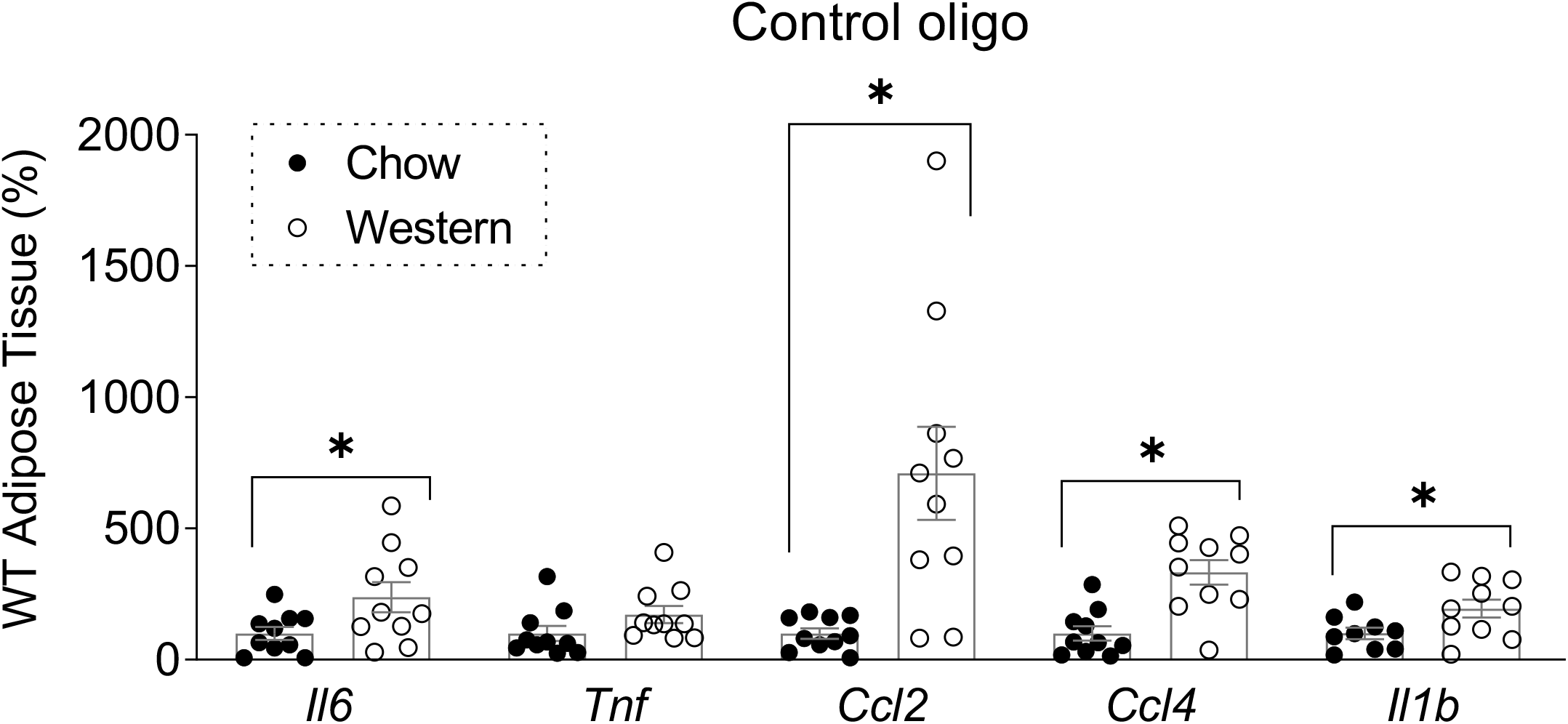
Adipose tissue inflammation in increased in WT mice on Western diet. Effect of Western diet on expression of pro-inflammatory markers (*Il6*, *Tnf*, *Ccl2*, *Ccl4*, *Il1b*) were assessed in whole adipose tissue by RT-qPCR from WT mice injected with control oligonucleotide. Data is normalized to β–actin. Data are relative to Chow ctrl diet average, n=10 mice/group. \**P*<.05

**Figure S5.**
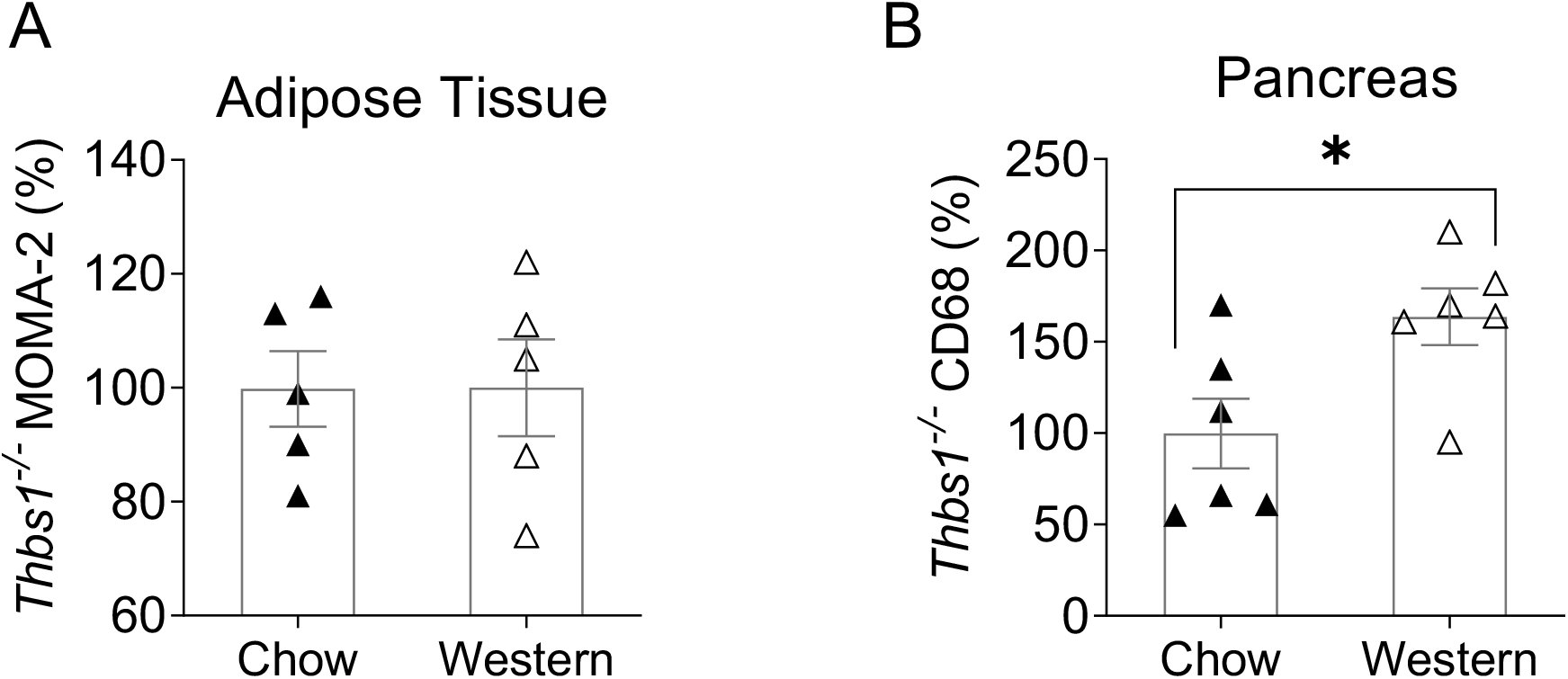
Effect of Western diet on macrophage accumulation in adipose tissue and pancreas in *Thbs1*^−/−^ mice. Effects of the Western diet on macrophage accumulation in adipose tissue and pancreas was determined by anti-MOMA-2 (A) or anti-CD68 (B) in *Thbs1*^−/−^ mice (injected with the control oligonucleotide), respectively. In AT, positive staining was normalized to mean adipocyte area for adipose tissue since adipocyte sizes were changed between groups. n=7 mice/group. Data are normalized to Chow ctrl diet average staining. \**P*<.05

**Figure S6.**
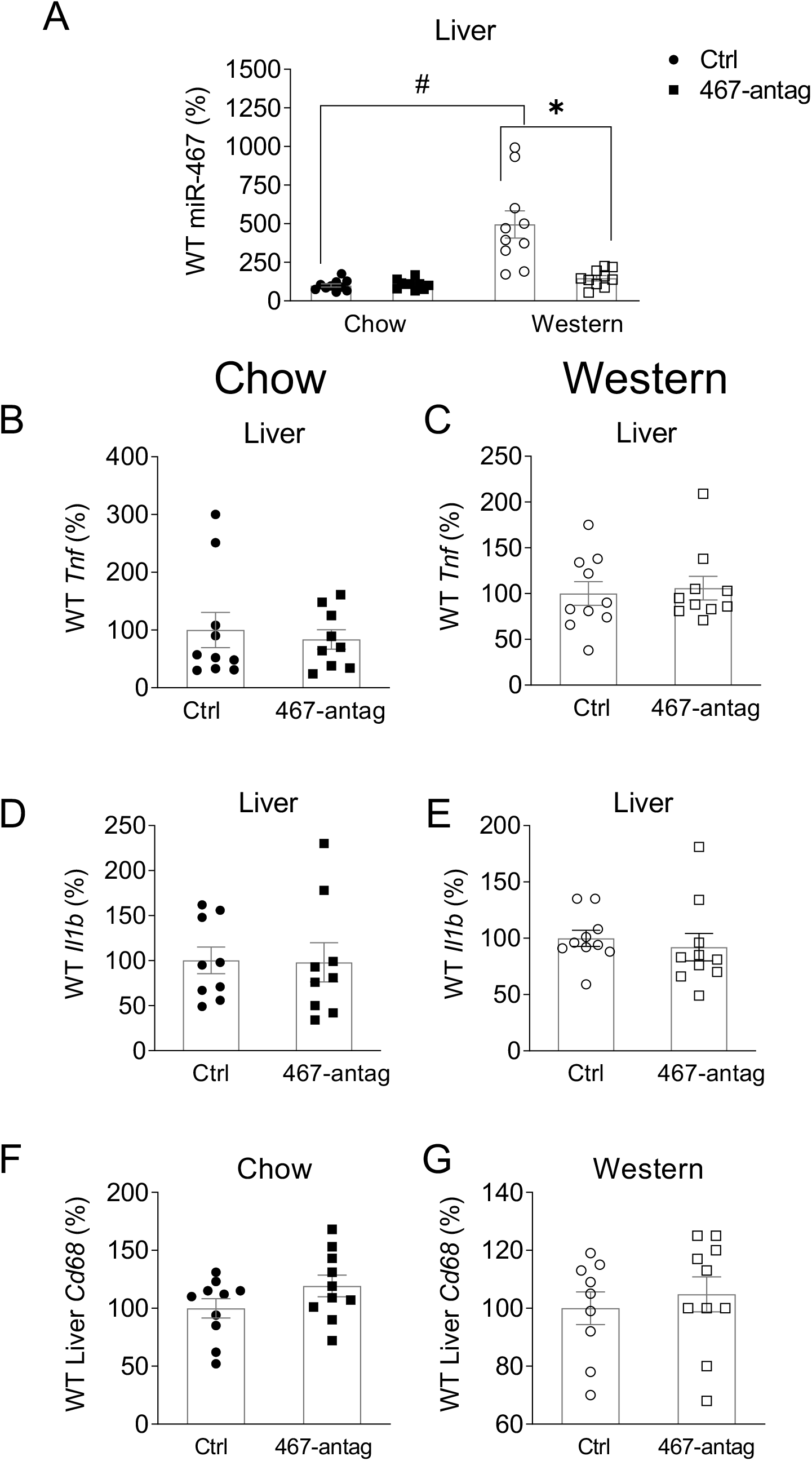
Effects of miR-467a-5p antagonist on liver. RNA from whole liver was extracted at the end of the experiment. Expression of miR-467 (A), inflammatory markers *Tnf* (B, C), *Il1b* (D, E), or macrophage marker *Cd68* (F, G) were assessed in WT mice on chow (B, D, F) or Western diet (C, E, G). Data are relative to ctrl oligo average (to Chow ctrl average in A). n=10 mice/group. \**P*<.05 vs ctrl oligo, # *P*<.05 vs chow diet

**Figure S7.**
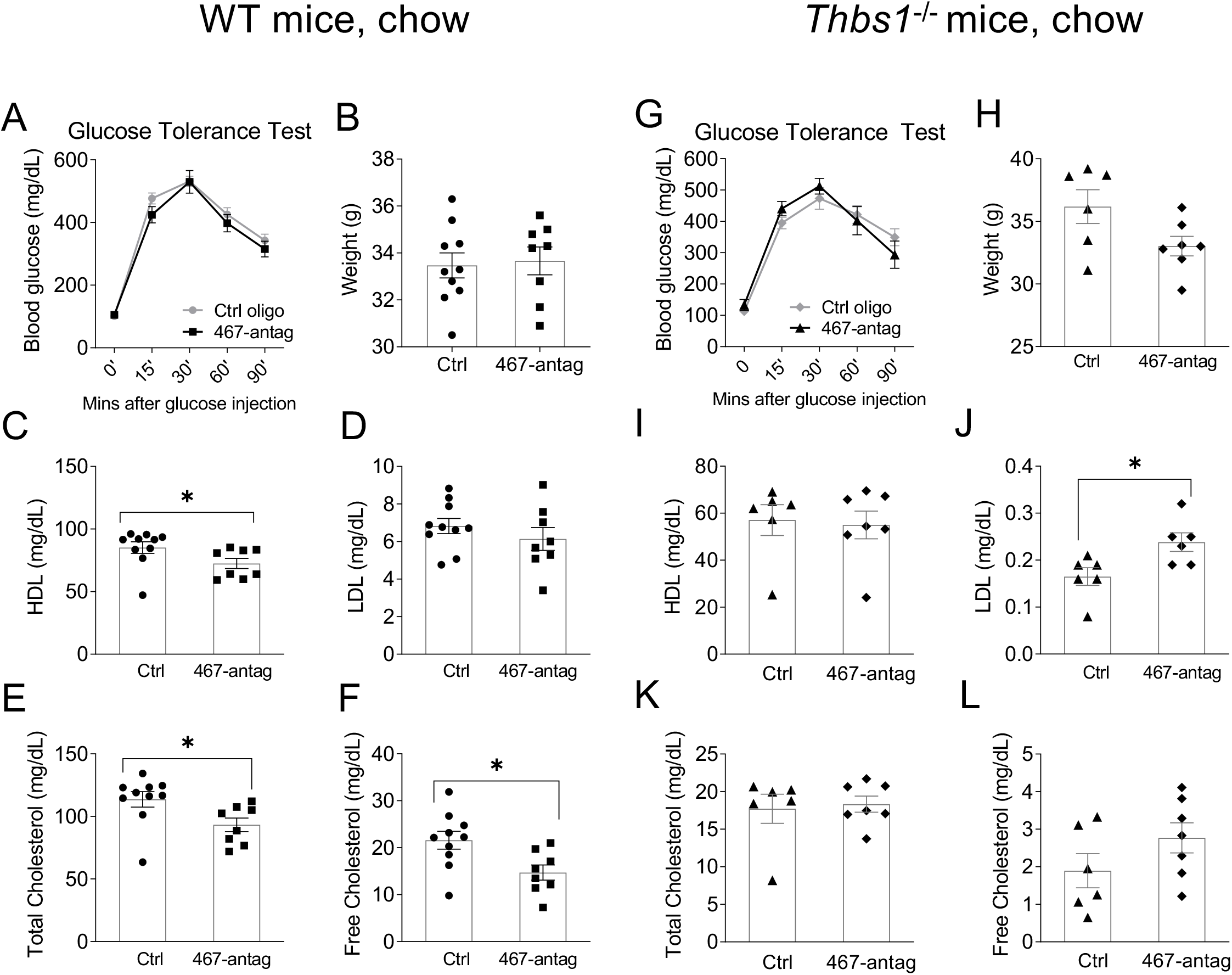
Effects of miR-467a-5p antagonist on mouse weight or blood lipid profile in chow-fed WT or *Thbs1*^−/−^ mice. Time course for the intraperitoneal glucose tolerance test (GTT) at the end of the experiment in chow-fed WT (A) or *Thbs1*^−/−^ (G) mice. Mouse weight measured at the end of study in WT (B) or *Thbs1*^−/−^ (H) mice. A quantification kit was used to quantify HDL (C, I), LDL (D, J), total cholesterol (E, K) or free cholesterol (F, L) from serum in WT or *Thbs1*^−/−^ mice, respectively. WT: n=10 mice/group. *Thbs1*^−/−^: n=7 mice/group. \**P*<.05

**Figure S8.**
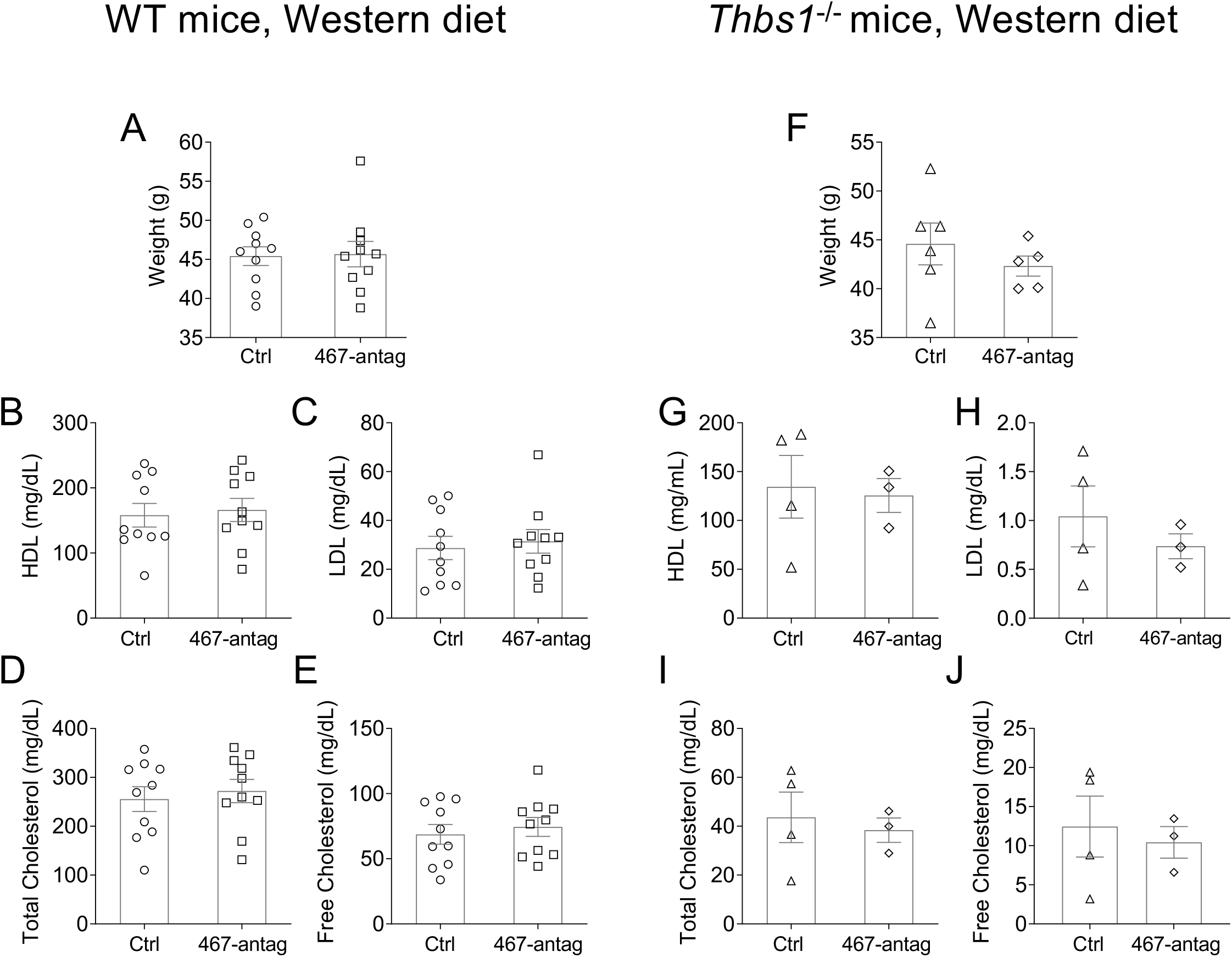
Effects of miR-467a-5p antagonist on mouse weight or blood lipid profile in WT or *Thbs1*^−/−^ mice on Western diet. Mouse weight measured at the end of study in WT (A) or *Thbs1*^−/−^ (F) mice on a Western diet. A quantification kit was used to quantify HDL (B, G), LDL (C, H), total cholesterol (D, I) or free cholesterol (E, J) from serum in WT or *Thbs1*^−/−^ mice, respectively. WT: n=10 mice/group. *Thbs1*^−/−^: n=7 mice/group.

**Figure S9.**
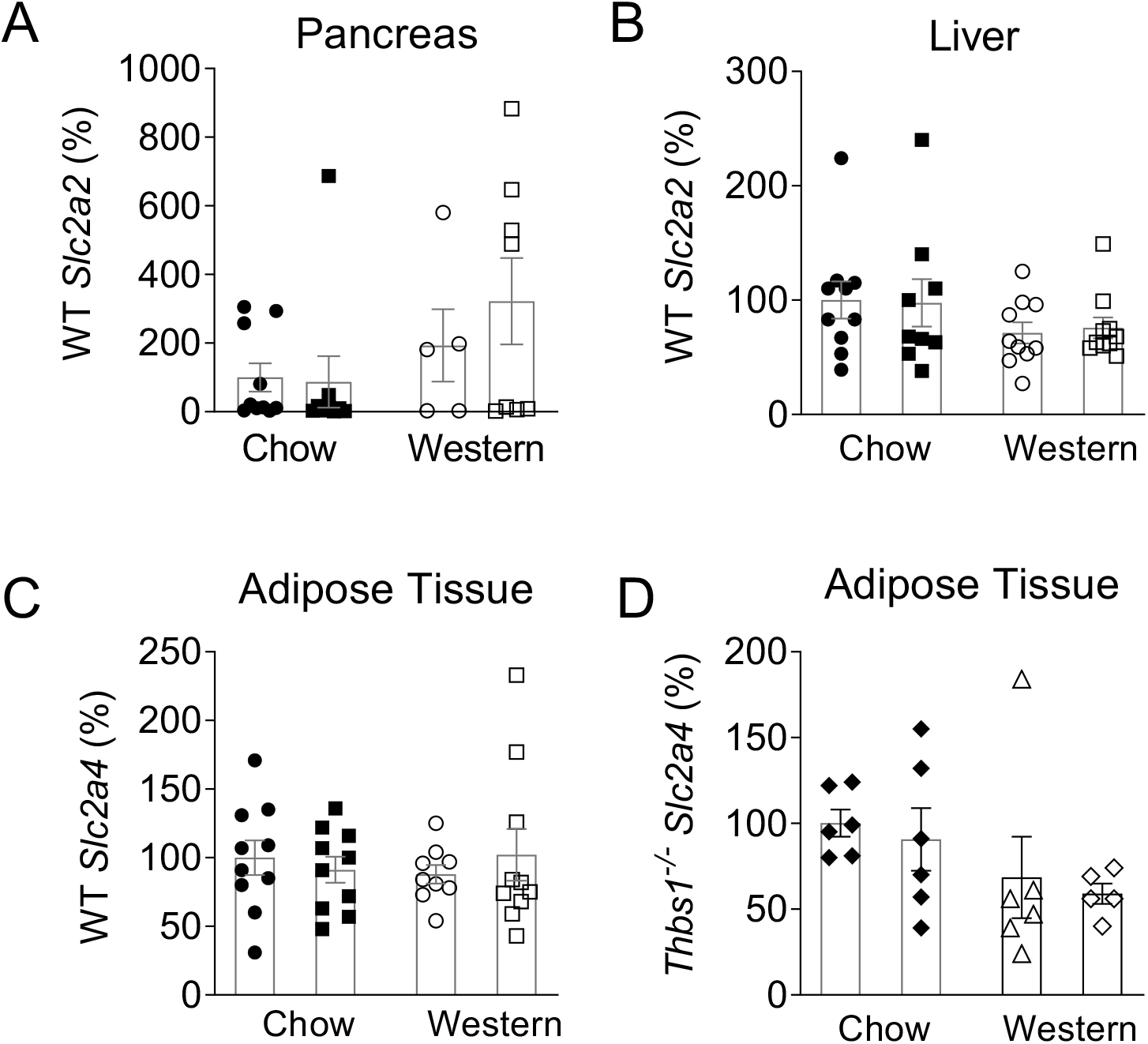
Expression of major glucose transporters in pancreas, liver, and adipose tissue. Expression of the major glucose transporters were measured: *Slc2a2* (Glut2) in pancreas (A) and liver (B) and *Slc2a4* in AT (C, D). Data are normalized to the chow ctrl average.

**Figure S10.**
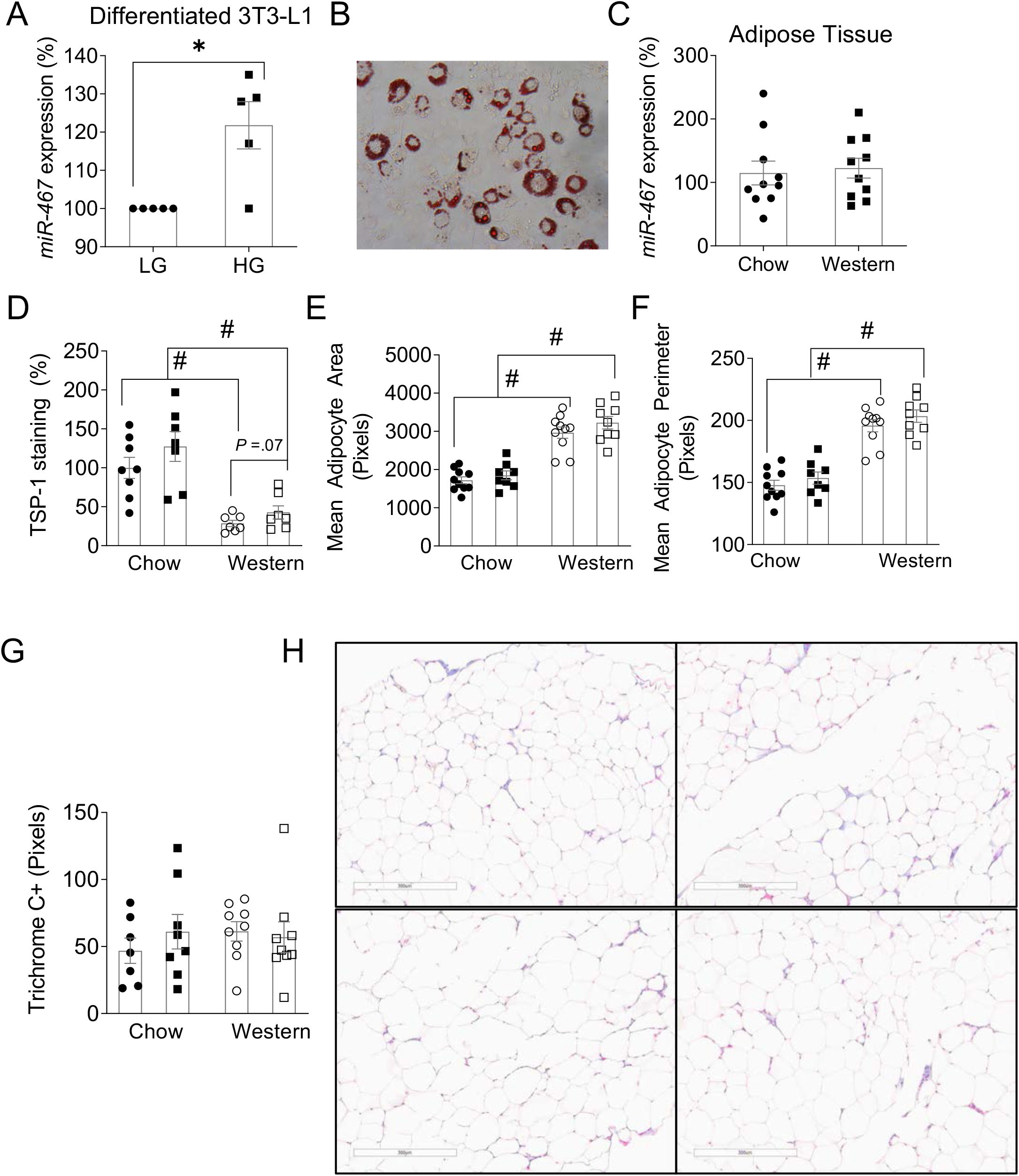
miR-467 in adipose tissue and the effects of the miR-467 antagonist injections. (A) miR-467 expression was measured 3 hrs post high glucose (HG) stimulation in cultured mouse fibroblasts (3T3-L1) differentiated into adipocytes. LG: low glucose control (5 mM D-glucose). HG: high glucose stimulated (30 mM D-glucose). Data is normalized to LG ctrl average. n=5 independent replicates. \**P* <.05. (B) Representative phase contrast image of Oil Red O Staining of 3T3-L1 cells at day 7 post-differentiation. 20x magnification. (C) Expression of miR-467 in WT C57/BL6 mouse adipose tissue on chow or Western diet for 32 weeks. n=10 mice/group. (D) Quantification of the % positive staining with an anti-TSP-1 antibody. (E) Mean adipocyte area and perimeter (F) were quantified from H&E-stained sections of adipose tissue. (G) Quantification of blue color density from Masson’s trichrome staining. (H) Representative images of trichrome staining. Scale bars at 300 µM. # *P*<.05 vs chow diet.

**Figure S11.**
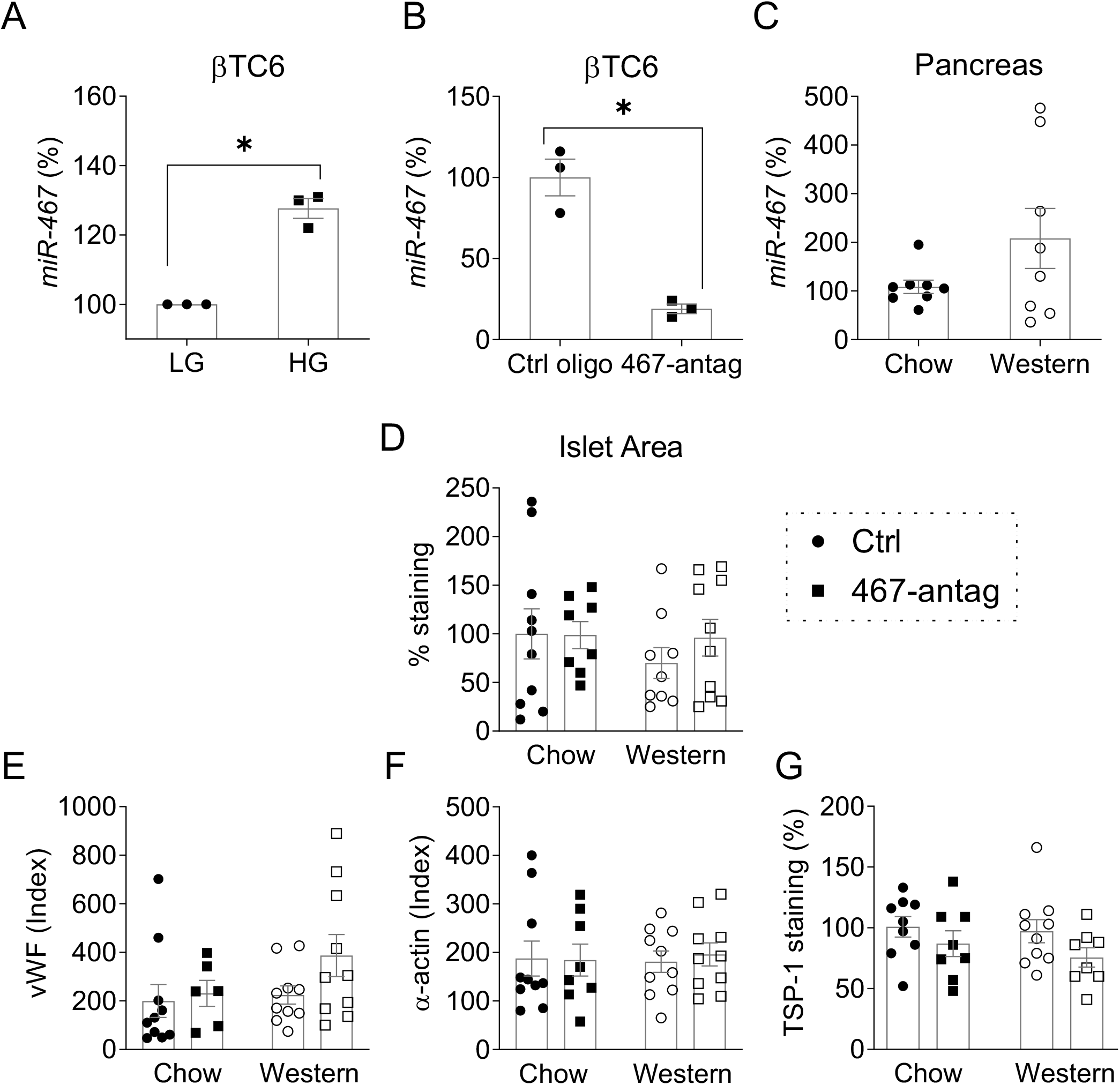
miR-467 in pancreas and the effects of the miR-467 antagonist injections. (A) miR-467 expression was measured 30’ post HG stimulation in a cultured mouse β cell line (βTC6). Data is normalized to LG ctrl average. n=3 independent replicates. LG: low glucose control (5 mM D-glucose). HG: high glucose stimulated (30 mM D-glucose). \**P*<.05 (B) Expression of miR-467 in C57/BL6 WT mouse pancreas on chow or Western diet for 32 weeks. n=10 mice/group. (C) Pancreas sections were stained for insulin and counterstained with hematoxylin. Islet area was quantified as % positive insulin staining over the total area per 100 pixels. Quantification of positive staining in pancreas using an anti-vWF (D), anti-α-actin, (E) or anti-TSP-1 (F) antibody.

**Figure S12.**
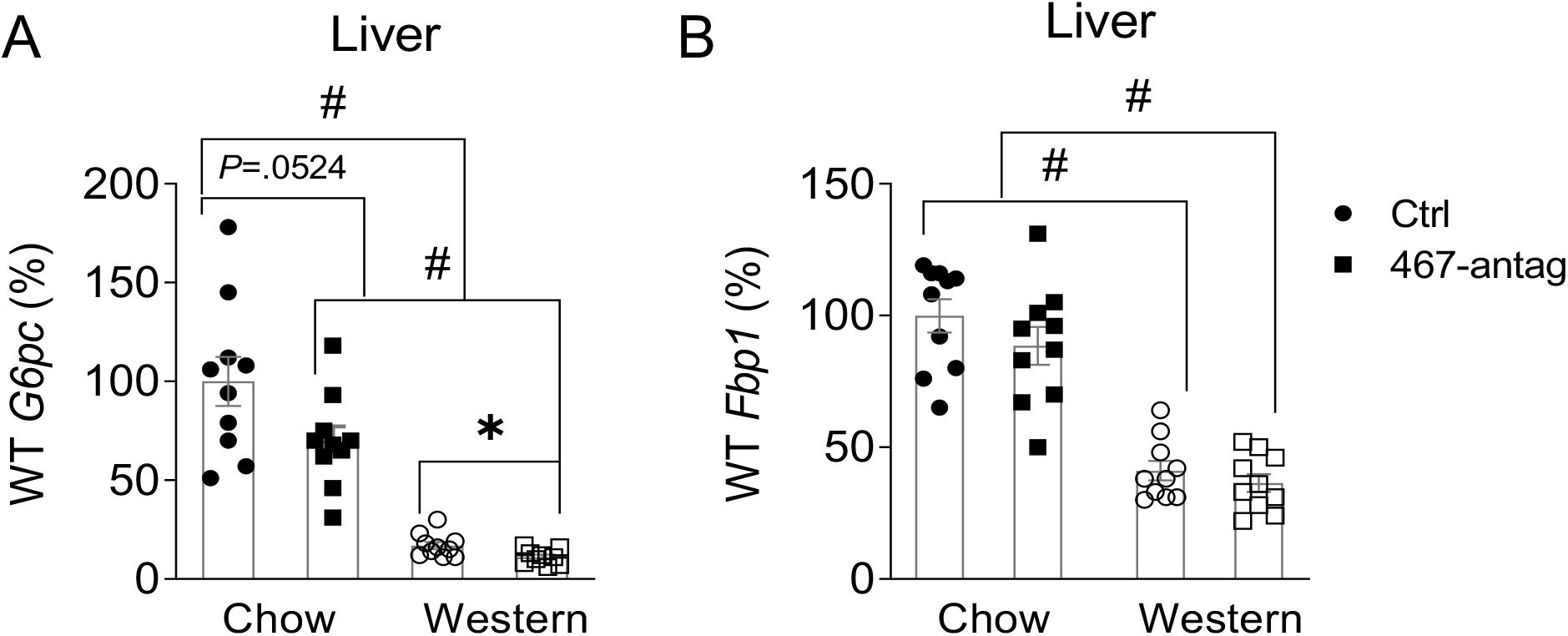
Gluconeogenesis genes in liver from WT mice. Expression of the gluconeogenesis genes, *G6pc* (A) or *Fbp1* (B) in liver were assessed in WT mice. Data are normalized to the chow ctrl average. n=10 mice/group. # *P*<.05 vs chow diet.

**Figure S13.**
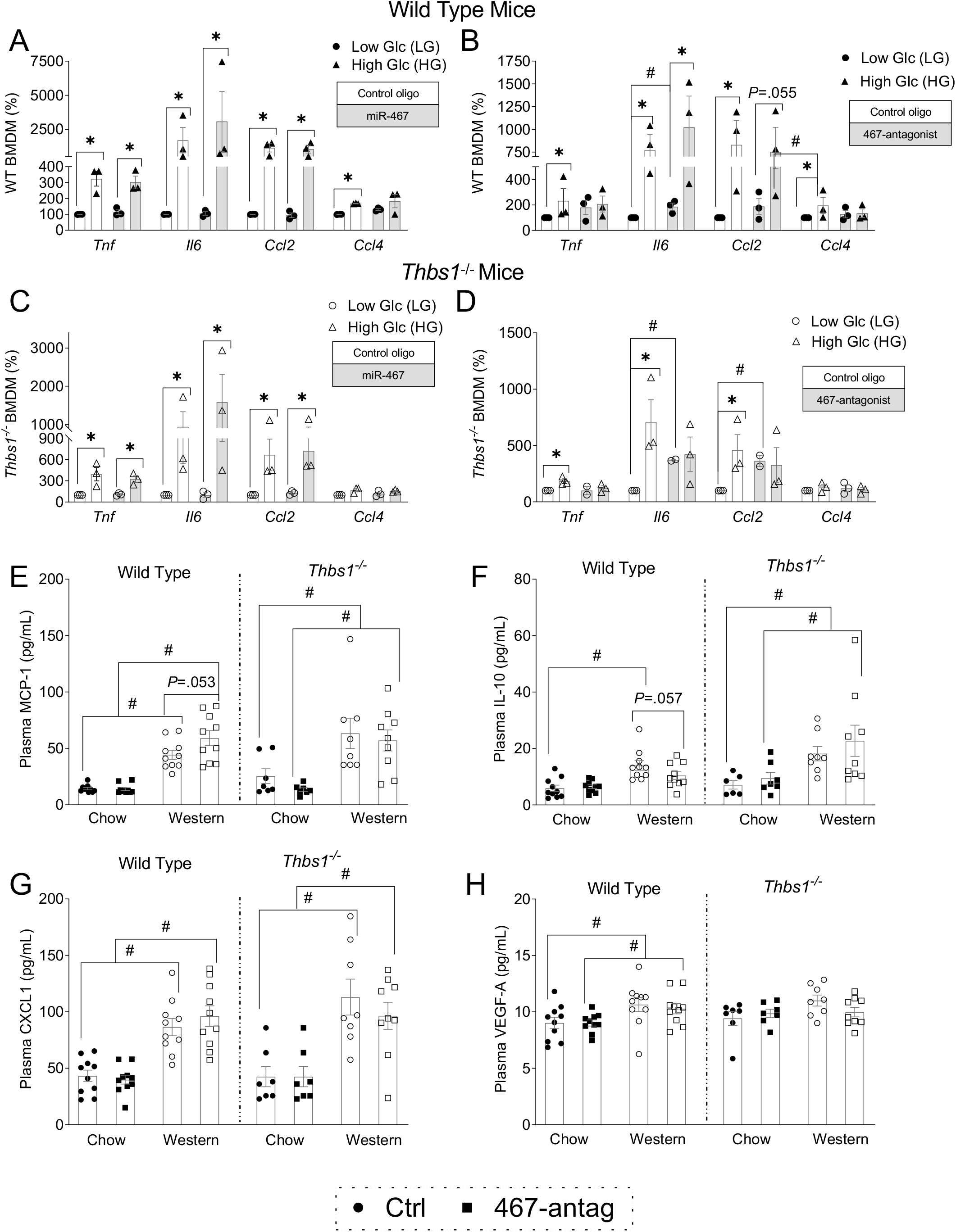
miR-467 blocks pro-inflammatory functions of cultured macrophages. Effect of miR-467 or miR-467 antagonist on expression of pro-inflammatory markers (*Tnf*, *Il6*, *Ccl2*, and *Ccl4*) were assessed in BMDM from (A, B) WT or (C, D) *Thbs1*^−/−^ mice transiently transfected with miR-467 (A, C) or a miR-467 antagonist (B, D) compared to a control oligo. RNA was collected 6 hrs post glucose stimulation by RT-qPCR and normalized to β–actin. Data are relative to the LG control stimulated samples per transfection, n=3 independent replicates. BMDM: bone marrow-derived macrophages. LG: low glucose control (5 mM D-glucose). HG: high glucose stimulated (30 mM D-glucose). \**P* <.05 compared to LG ctrl. # *P* <.05 vs chow diet. (E – H) Plasma from WT or *Thbs1^−/−^* mice, collected at 32 weeks, were assayed by U-plex for circulating (E) MCP-1, (F) IL-10, (G) CXCL1 and (H) VEGF-A. # *P* <.05 vs chow diet.

